# Proteoform plasticity modulates the temperature response in circadian timekeeping

**DOI:** 10.64898/2026.09.05.749499

**Authors:** Jacqueline F. Pelham, Alex T. Keeley, Emery T. Usher, Jacob H. Martinsen, Meaghan S. Jankowski, David Moses, Alexander E. Mosier, Joshua Thomas, Nathan D. Buckley, Birthe B. Kragelund, Shahar Sukenik, Alex S. Holehouse, Jennifer M. Hurley

## Abstract

The molecular circadian clock optimally coordinates an organism’s physiology with the light/dark cycle. One commonality among circadian molecular clock proteins is that they undergo alternative splicing, yielding multiple proteoforms. The isoform-specific sequences spliced into clock proteins contain intrinsically disordered regions (IDRs), suggesting that these regions might be intricately involved in cellular regulation. However, the functions of these isoform-specific IDRs in the clock remain poorly defined. Here, we use the core clock repressor FREQUENCY (FRQ) from the fungal circadian model system *Neurospora crassa* to test the hypothesis that alternative splicing of core clock IDRs yields multiple proteoforms that expand the functional regulatory capacity of FRQ. To do so, we biophysically characterized an IDR specific to the long isoform of the core clock repressor FRQ and identified motifs and molecular behaviors that regulate clock robustness in a temperature-dependent manner. We further compared the interactomes of two FRQ isoforms, identifying distinct functions and interacting proteins that could contribute to temperature regulation. Taken together, our data highlight that isoform-specific IDRs in clock-repressor proteins enhance and expand clock function in a context-dependent manner.

**Significance Statement:** Circadian clocks align internal physiology with daily environmental cycles across diverse life forms, yet the molecular mechanisms that confer robustness under fluctuating conditions remain poorly understood. Here, we show that intrinsically disordered regions (IDRs) within isoform-specific variants of the clock protein FREQUENCY encode temperature-responsive regulation and timing. We identify a motif that shapes conformational behavior and, when altered, selectively disrupts timekeeping in a temperature-dependent manner. Our findings establish a link among IDR sequence, conformational dynamics, and organismal phenotype and support a model in which IDRs function as environmental rheostats. These mechanisms likely extend to proteins beyond circadian systems.

## Introduction

The regular environmental changes that accompany the oscillatory day/night cycle on Earth have prompted the widespread evolution of a molecular circadian clock that tunes organismal physiology to this 24-hour cycle (1). These clocks broadly regulate rhythms to enhance an organism’s competitiveness in its environment (1–4). Disruption of circadian rhythms can lower fitness and increase disease risk (5, 6). In eukaryotes, the circadian clock is an architecturally conserved transcription-translation feedback loop (TTFL). In fungi and animals, the TTFL is orchestrated by activating and repressing proteins that coordinate with auxiliary feedback loops to regulate circadian physiology (Fig. 1a) (7). Although the proteins involved in the TTFL are mostly defined, much remains to be discovered about the molecular determinants and biophysical mechanisms that enable robust circadian timekeeping within this TTFL.

**Figure 1.**
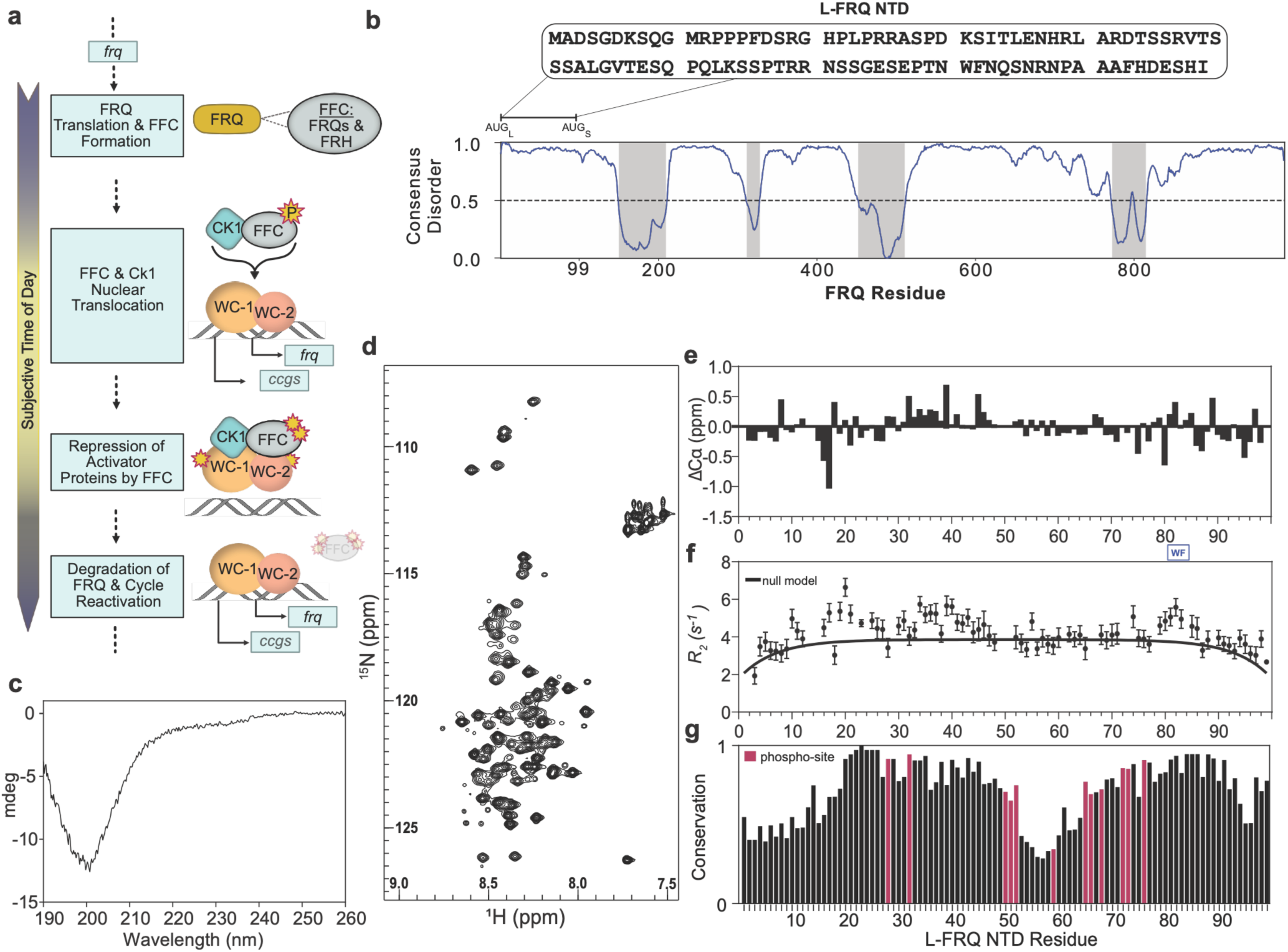
The circadian repressor FRQ has an intrinsically disordered N-terminal domain (NTD) specific to its long isoform. **a**. The circadian cycle is orchestrated via a protein-based, temporally regulated, transcription- translational feedback loop (TTFL). Late in the subjective night, the repressor *frq* is translated and binds to its partner protein FREQUENCY-interacting RNA helicase (FRH), forming the FRQ-FRH Complex (FFC). The FFC binds with Casein Kinase-1 (CK1) and is translocated to the nucleus, where it represses the heterodimer transcriptional activators White Collar-1 and -2 (WC-1 and WC-2, referred to as the White-Collar Complex, WCC).

One conserved property of clock circuitry that enables robust timekeeping is the ability to respond to and buffer temperature fluctuations. This includes being temperature-compensated (maintaining a constant period despite altered kinetics due to temperature changes) and being plastic to temperature changes (distinguishing relevant temperature cues, such as seasonal cycles, from temperature changes that require buffering) (8, 9). While the precise mechanisms underlying these temperature-related phenomena remain to be determined, it has been proposed that temperature-induced alternative splicing (AS) mediates the temperature responses of the TTFL (9–12). In animal and fungal clocks, temperature- induced AS of core clock repressors yields multiple proteoforms, each playing a unique role in temperature responses and adaptation of the clock (9–14).

In fungal and animal clocks, temperature-dependent AS results in the addition or omission of N- or C-terminal regions predicted to be intrinsically disordered in the TTFL repressor proteins (7). This is relevant because intrinsically disordered regions (IDRs), which exist as ensembles of interconverting conformations, exploit these ensembles to act as sensors influenced by environmental and context-specific factors (e.g., temperature, pH, and post-translational modifications (PTMs)) (7, 15, 16). In line with this, IDRs expand a protein’s potential to participate in a wider range of biological functions and processes in a context-dependent manner (15, 17, 18). Considering the tunability of IDRs and the temperature-dependent AS of these regions, we hypothesized that the clock extends its functional range and capabilities by incorporating (or removing) temperature-specific IDRs, thereby leveraging IDR-specific proteoform plasticity to facilitate the temperature response.

To test our hypothesis, we characterized the isoform-specific region of the circadian repressor FREQUENCY (FRQ) in the model organism *Neurospora crassa.* FRQ undergoes AS, resulting in the incorporation or omission of a 99-amino-acid N-terminal domain (NTD). The isoform with the NTD is referred to as Long-FRQ (L-FRQ, 989 amino acids) (Fig. 1b) and is dominant and up-regulated at ambient and warmer temperatures, respectively. As temperatures decrease, L-FRQ expression decreases, and the Short-FRQ isoform (S-FRQ, 890 amino acids), lacking the NTD, is increased (13). We demonstrated that the L-FRQ NTD is intrinsically disordered and exhibits ensemble-affecting, sequence-specific features, including increased solvent accessibility. We further identified sequence-specific motifs that, *in vivo*, affect clock robustness and proper period maintenance in a temperature-dependent manner, indicating that the isoforms have distinct half-lives and interact with different partners. Finally, we found discrete biochemical features and differences in clock-regulatory capacity between S- and L-FRQ. Taken together, our data suggest that proteoform-specific functions in the TTFL are in part facilitated by the disordered sequence features of the L-FRQ NTD, and that the NTD can enhance clock timekeeping and robustness in a temperature-dependent manner. Given the enrichment of disorder in circadian clock proteins in fungal and animal clocks (7, 19), this work provides mechanistic insights into how the AS of IDRs may enable and expand clock regulation, particularly in the context of the clock’s temperature response.

## Results

### The L-FRQ N-terminal domain is disordered and has conserved, dynamically distinct regions

Given the tractability of FRQ and *N. crassa* as a model for IDR-associated circadian phenomena, we sought to explore the ensemble features of its temperature-regulated NTD to identify a mechanism for proteoform plasticity. We hypothesized that specific sequence features of FRQ associated with protein disorder and phosphorylation could facilitate isoform-specific regulation. To test this hypothesis, we first computationally explored the ensemble characteristics of the region unique to the long FRQ isoform. Consistent with previous reports, general analysis by Metapredict suggested that the isoform-specific N-terminal domain (NTD) of L-FRQ (residues 1-99) was predicted to be disordered (Fig. 1b) (19–21). To experimentally validate the predicted disordered nature of the L-FRQ NTD, we expressed and purified the L-FRQ NTD protein fragment (residues 1-99, see Methods) and analyzed it in parallel by circular dichroism (CD) and NMR spectroscopy.

The CD spectrum of L-FRQ NTD showed a deep negative minimum around 200 nm, which is congruent with the characteristics of an IDR (Fig. 1c). These data were corroborated by the HSQC spectrum of the L-FRQ NTD, which showed a narrow peak dispersion in the ^1^H dimension, further indicating the disordered nature of the L-FRQ NTD (Fig. 1d). Following NMR assignments, the small values in the secondary structure profile calculated from the secondary Cα chemical shifts (ΔCα) further support a lack of stable secondary structure in the L-FRQ NTD (Fig. 1e). Some consecutive negative (around F16-S18, N86-D95) and positive values (D30-T49) indicate the presence of lowly populated extended and helical structures, respectively. Furthermore, we observed a small set of low-intensity NMR peaks related to proline *cis-trans* isomerization for Pro23 (22, 23). Interestingly, the per-residue transverse relaxation rates (*R*_2_) show subregions in the L-FRQ NTD that have slower local dynamics compared to a random coil model (Fig. 1f). Notably, the regions with slower local dynamics, which we will refer to as “dynamically distinct”, overlap with the regions of the NTD with lowly populated preferred structures and are retained within the regions with highest amino acid conservation (Fig. 1f-g). The presence of these dynamically distinct regions is likely related to the presence of transient contacts and suggests the L-FRQ NTD ensemble is structurally biased.

The WCC regulates the expression of both *frq* and many other genes not involved in the core clock mechanism, referred to as clock-controlled genes (*ccgs*). CK1 and other kinases progressively phosphorylate (yellow stars) the WCC and FRQ to progress the cycle. Once FRQ is hyperphosphorylated, it can no longer repress the WCC and is targeted for degradation, allowing the cycle to begin again. **b.** The sequence of the L-FRQ NTD is highlighted above the predicted protein disorder plot of the FRQ sequence, generated by Metapredict (24). The transcriptional start sites for each of the FRQ isoforms are noted as AUG_L_ (the long isoform or L-FRQ) and AUG_S_ (the short isoform or S-FRQ). Splicing in the upstream regions of *frq* transcripts results in two nearly identical isoforms of FRQ, with the only difference being the L-FRQ N-terminal domain (NTD), which spans residues 1-99 in L-FRQ. Greyshaded areas indicate regions with predicted structure (<0.5 Metapredict score). **c.** Far-UV circular dichroism (CD) spectra of the L-FRQ NTD. **d.** ^1^H, ^15^N HSQC spectrum of the L-FRQ NTD recorded at pH 7.4, 10 °C. **e.** Secondary Cα chemical shifts for the L-FRQ NTD. **f.** Transverse relaxation rates (*R*_2_) for the L-FRQ NTD. The line corresponds to *R*_2_ values of a random coil (25), and error bars are standard errors from fits. **g.** Weighted conservation for the L-FRQ NTD from FRQ homologs. Maroon bars indicate amino acids that are phospho-sites as identified by (26, 27).

### Sequence-biased regions in L-FRQ NTD are tuned by phosphorylation and a conserved aromatic pair

Given the conservation of the dynamically distinct regions, we hypothesized that these regions may have context-specific chemical signatures that can influence ensemble behavior. To investigate the contribution of conserved sequence context to the local and global ensemble behavior of the L-FRQ NTD, we developed a computational method (FINCHES-DMS) to perform exhaustive deep mutational scanning (DMS) to assess the sequence-dependence of predicted IDR-mediated interactions.

FINCHES-DMS builds on FINCHES, a computational approach that repurposes the chemical physics developed for coarse-grained force fields to determine the interaction potential between pairs of sequences (28). In FINCHES-DMS, we provide either a single sequence (as here, for intramolecular interactions) or a pair of sequences (for intermolecular interactions). Each position in the sequence is systematically mutated to every other possible amino acid, and for each single-point mutation, the predicted interaction between it and either its own (intra) or a partner sequence (inter) is recomputed (see *Methods*). While FINCHES-DMS is limited to interactions driven by chemical specificity, it allows us to rank order residues in a sequence to identify those most strongly predicted to impact intra- (or inter-) molecular interactions for further biophysical or functional investigation.

Using FINCHES-DMS, we identified multiple aromatic residues that, when mutated, impacted the predicted intramolecular interactions of the NTD. The most outsized impact stemmed from a pair of neighboring residues, 81 (W) and 82 (F) (Fig. 2a). Intriguingly, the identified aromatic pair (which going forward we will refer to as the “WF pair”) occurs within one of the dynamically distinct regions of the FRQ NTD we observed by NMR (Fig. 1f). Considering the well-known role of aromatic residues serving as “stickers” promoting attractive interactions, we speculated that the WF pair may tune intra or intermolecular interactions within the L-FRQ NTD (29).

**Figure 2.**
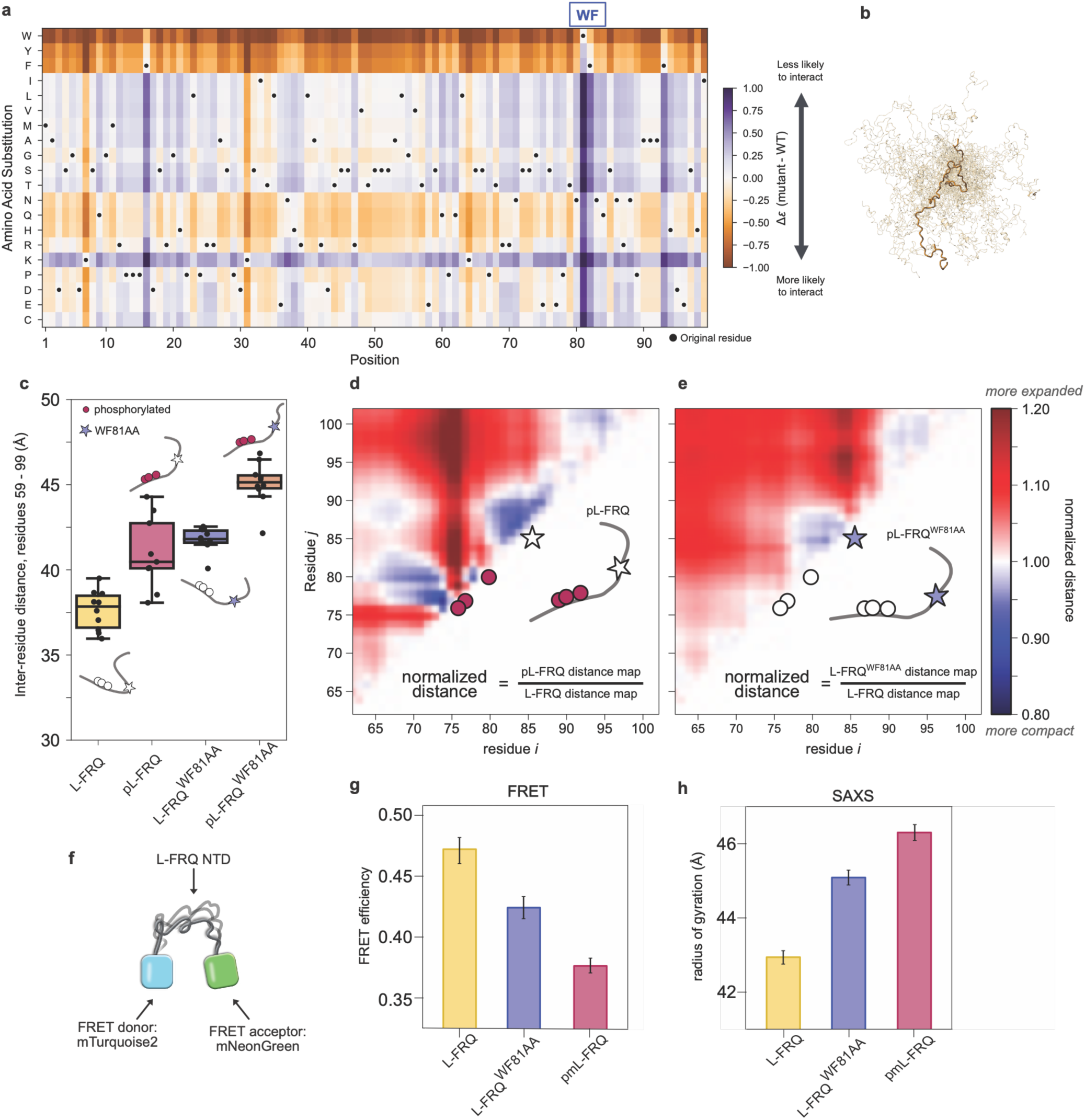
The L-FRQ NTD exhibits ensemble biases influenced by phosphorylation and aromatic residues. **a**. A heat map of a deep mutational scan via coarse-grained simulations (FINCHES-DMS) depicts intermolecular interaction potentials across the L-FRQ NTD. **b.** Exemplars of the superimposed all-atom Monte-Carlo simulations of the L-FRQ NTD. **c.** Inter-residue distances were calculated from all-atom Monte Carlo simulations of the phosphorylated (pL-FRQ) NTD and mutant NTDs. The associated diagrams serve as models of the simulation outputs. **d-e.** Contact maps comparing the interchain contact frequency based on the Monte-Carlo simulations of pL-FRQ NTD residues (**d.**) and WF81AA mutant (**e.**), normalized by the native sequence (L-FRQ NTD). **f.** A schematic of the SAXS and ensemble FRET (eFRET) constructs. The L-FRQ NTD is flanked by mTurquoise2 and mNeonGreen at the N- and C-termini, respectively. **g.** The calculated eFRET efficiency for the L-FRQ NTD constructs. Error bars show the spread of data from 12 replicates over two repeats. The phosphomimetic variant is indicated by pmL-FRQ. **h.** The radii of gyration of the eFRET constructs were determined by Small Angle X-ray Scattering (SAXS).

Notably, within a few residues of the conserved WF pair, there are three validated phospho-sites that are temporally phosphorylated and known to affect the circadian period when mutated (S72A, S73A, and S76A), which we refer to as the “ phospho-cluster “ (Fig. 1g) (26). Considering the proximity of the phospho-cluster to the WF pair, we hypothesized that the temporal phosphorylation of this cluster may impact local ensemble behavior in conjunction with this aromatic pair (30). To evaluate the ability of phosphorylation to combine with the WF pair to induce changes in the ensemble of the L-FRQ NTD, we turned to all-atom Monte Carlo simulations (Fig. 2b). We iterated various models of the NTD, including a model in which the WF residues are mutated to an alanine pair (AA) (L-FRQ^WF81AA^). Phosphorylation of the phospho-cluster was also incorporated (pL-FRQ), along with the combination of these two sequence modifications (pL-FRQ^WF81AA^).

Comparing the inter-residue distance of residues 59-100 among all these models, we found that all modified sequences were more expanded than the native sequence (L-FRQ) (Fig. 2c). The phosphorylated version of the NTD (pL-FRQ) was more expanded than the native sequence, likely due to the additional two negative charges, increasing the energetic favorability of solvent exposure. When comparing the contact frequency of pL-FRQ to L-FRQ, we found both repulsion and expansion occur because of phosphorylation (Fig. 2d). Meanwhile, the dimensions of pL-FRQ were not statistically different from the model of L-FRQ^WF81AA^ (p-val 0.270, Table S1). Likely due to the chemistry of the aromatic pair, the L-FRQ^WF81AA^ iteration also induced expansion at the C-terminus differentially as compared to pL- FRQ. However, the largest end-to-end distance was observed in pL-FRQ^WF81AA^, suggesting that each of the two regions contributes independently to the local context of this region (Fig. 2c). These findings collectively indicate that the region undergoes progressive solvent exposure during temporal phosphorylation and that the WF pair plays a parallel role in this expansion.

To confirm our *in silico* findings *in vitro* and test our hypothesis that the WF pair impacts the ensemble of the L-FRQ NTD, we next measured the ensemble behavior of purified L-FRQ NTD using Size Exclusion Chromatography Small Angle X-ray Scattering (SEC-SAXS) and ensemble Förster Resonance Energy Transfer (eFRET). Briefly, for eFRET, the IDR of interest is placed between two fluorescent proteins, and the resultant FRET efficiency serves as a proxy reporting on the ensemble dimensions (31) (Fig. 2f). We purified FRET constructs in which one of the versions of interest of the L-FRQ NTD was inserted between donor (mTurquoise2) and acceptor (mNeonGreen) fluorescent proteins. Compared with the wild-type L-FRQ NTD, we observed decreased eFRET efficiency (ensemble expansion) in mutants where the WF was altered (L-FRQ^WF81AA^). Similarly, a phosphomimetic variant (pmL-FRQ) mirroring the phosphorylation pattern observed in pL-FRQ also reduces eFRET efficiency. These results are fully consistent with our *in silico* results, suggesting that the WF mediates intramolecular interactions mostly with other hydrophobic residues (which are weakened in the WF81AA mutant) and that negative charges from phosphorylation/phosphomimetic mutations introduce electrostatic repulsion that mediates expansion (Fig. 2g). We next corroborated the eFRET data with SEC-SAXS using the purified protein from the constructs described above. All L-FRQ NTD constructs produced clean single peaks, indicating monodisperse samples (i.e., free of oligomerization and aggregation) (Fig. S1a). When comparing the resultant SEC SAXS chromatograms and the SAXS-calculated radius of gyration (*R_g_*) for the native L-FRQ NTD (L-FRQ), L-FRQ^WF81AA^, and pL-FRQ, we observed an expansion effect of the WF81AA and phosphomimetic mutations, which correlated with our *in silico* calculations (Fig. 2g-h and Fig. S1a-b).

In conclusion, FINCHES-DMS, simulations, eFRET, and SEC-SAXS all paint a consistent picture: the WF facilitates attractive intramolecular interactions, which are reduced upon mutation to AA, and phosphorylation drives chain expansion likely through electrostatic repulsion (Fig. 2). These data support the notion that dynamically distinct regions, encoded by sequence specificity within the L-FRQ NTD (i.e., the WF pair), tune the ensemble behavior of the L-FRQ NTD. With this in mind, we next wondered how changes in NTD ensemble might relate to circadian function.

### A dynamically distinct region of the FRQ NTD affects circadian periodicity

Given their effects on the L-FRQ NTD ensemble, we hypothesized that mutations within and around the dynamically distinct regions described above would affect FRQ’s ability to regulate circadian timing *in vivo*. To directly test this hypothesis, we created strains of Neurospora that were mutated at the WF pair and phospho-cluster sites in FRQ (Fig. 3a). To do so, we targeted mutant alleles of FRQ with a C-terminal epitope tag (V5, 10-His, 3-Flag) to the cyclosporin (*csr-1*) locus of a *frq* KO strain that displays no circadian patterns (32) (Fig. 3a and 3b). This FRQ KO strain also harbors a reporter construct in which a minimal *frq* promoter is fused to a luciferase coding gene at an exogenous locus (33, 34). Thus, using these strains, we were able to track luciferase expression as a proxy for the molecular clock period *in vivo*. To control for the possibility that the short isoform of FRQ might mask the phenotypes from the FRQ alleles, we altered the FRQ start codon in these alleles to prevent translation, yielding L-FRQ-only strains (Fig. 3a) (see Methods for strain details). For both the phospho-cluster and WF pair strains, we substituted the key residues with Ala (S72A, S73A, and S76A, or W81A and F82A, referred to as strain L-FRQ^3pS/3A^ and L-FRQ^WF81AA^, respectively) (Fig. 3a).

**Figure 3.**
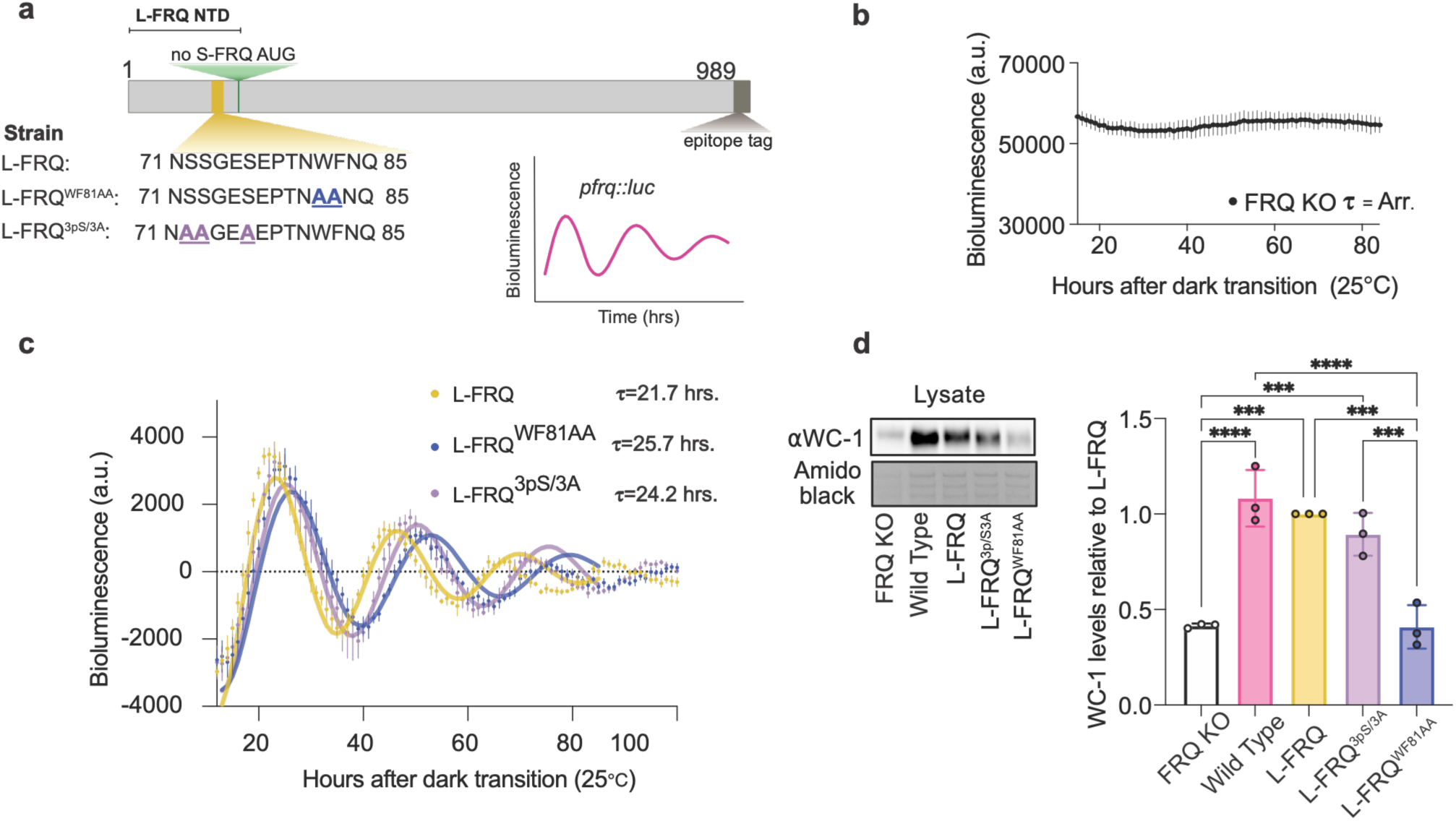
L-FRQ NTD mutants in the dynamically distinct WF pair region affect timekeeping, with only the WF pair mutant altering the levels of the circadian activator WC-1. **a**. A schematic of the details of the mutations designed in the FRQ NTD to test the function of the dynamically distinct regions. Constructs were designed to alter the S-FRQ start codon and mutate the key residues of interest in the L-FRQ NTD. The C-terminus of each construct included an epitope tag (10xGly, V5, 6xHis, 3FLAG). All the constructs were transformed into a *frq* KO strain containing the minimal C-box *frq* promoter fused to the luciferase gene (*pfrq::luc*). b. The smoothed, normalized, and detrended bioluminescence trace of the FRQ-KO strain into which the L-FRQ constructs were transformed. Points represent the average of n=6 wells, with bars indicating the standard deviation at each time point. c. The smoothed, normalized, and detrended luciferase reporter activity of the L-FRQ NTD mutant strains. Points represent the average of n=6 wells, with bars indicating the standard deviation at each time point. The solid trace represents the fit determined by the Chronostar algorithm used to calculate the corresponding period (τ) of each strain (35) Periods differ significantly across all three strains and are plotted in Fig S2a. d. Densitometry analysis of the relative level of WC-1 in the L-FRQ NTD mutants relative to the L-FRQ strain. Western blot is a representative of n = 3 biological replicates, which were normalized to an amido black stain for each replicate. Relative levels of WC-1 were tested with a One-way ANOVA (F = 23.09, p<0.0001) and Tukey’s multiple comparisons (***p<0.001, **** p<0.0001).

Luciferase analysis of these strains revealed that at 25°C, the phospho-cluster mutant (L-FRQ^3pS/3A^) and the WF pair mutant (L-FRQ^WF81AA^) both displayed longer periods than the L-FRQ control (L-FRQ^WF81AA^ τ=25.7hrs and L-FRQ^3pS/3A^ τ=24.2hrs, vs L-FRQ τ=21.7 hrs) (Fig. 3c). The period of the L-FRQ^3pS/3A^ strain agrees with previous reports of an Ala substitution of the phospho-cluster by Baker et al. (26). However, unlike the previous findings, the L-FRQ^3pS/3A^ strain did not have a lower amplitude rhythm compared to the L-FRQ control (Fig. S2b) (26). In this assay, the period reports on the length of one circadian cycle, while the amplitude reports on the amount of *frq* activation by WC-1. While the phospho-cluster (L-FRQ^3pS/3A^) strain had a similar amplitude to the control, the L-FRQ^WF81AA^ mutant had a statistically significantly higher amplitude, suggesting an additional mechanism may be at play in the WF mutant compared to the phospho-cluster strain, resulting in the differential phenotypes (Fig. S2b). Taken together, these data demonstrate that L-FRQ NTD mutations in a dynamically distinct region affect the macromolecular mechanics of circadian timekeeping, supporting our hypothesis that these regions modulate FRQ’s circadian function.

We next asked if our L-FRQ^WF81AA^ and L-FRQ^3pS/3A^ mutations affect clock function by regulating the expression and interactions of other core clock proteins, as has been shown for other FRQ regions (32, 36, 37). To investigate changes in core clock protein levels, we performed a semi-quantitative Western blot analysis on lysates of the L-FRQ^WF81AA^, L-FRQ^3pS/3A^, and L-FRQ strains. We first compared total FRQ protein levels in the L-FRQ strain (i.e., a strain where only L-FRQ is expressed) to total FRQ protein levels in our NTD mutants. We found that FRQ protein levels in the L-FRQ^3pS/3A^ strain were not statistically different from the L-FRQ control strain, while the FRQ^WF81AA^ strain had roughly half the amount of FRQ as the L-FRQ strain (Fig. S3a-b). However, when we performed the same analysis on a strain that expressed both the S and L-FRQ isoform (as would be seen in nature) with the same epitope tags and recombineering strategy (referred to as the dual isoform strain (FRQ^D^) (*csr-1::pfrq::frq^v5his103flag^*), we found that levels of FRQ protein were similar in the L-FRQ^WF81AA^ and FRQ^D^ strains (Fig. S3a-b). Furthermore, the FRQ^D^ strain exhibits a period phenotype that closely resembles the L-FRQ strain τ=21.9 hours (Fig. S4b-c). These data suggest that changes in FRQ protein levels in the mutants are not responsible for the long-period phenotype in the L-FRQ^WF81AA^ strain.

To investigate the molecular origins of the L-FRQ^WF81AA^ phenotype, we surveyed the levels of the core clock proteins WC-1 and FRH. WC-1 is one of the heterodimeric transcription factors that activates *frq* expression, while FRH is part of the repressive complex of the clock, serving as a ‘nanny’ for FRQ (20) When we examined the levels of clock proteins in the L-FRQ^3pS/3A^ strain relative to the L-FRQ strain, we found no difference for FRQ, FRH, or WC-1 levels (Fig. 3d and Fig. S3). However, for the L-FRQ^WF81AA^ strain, we found a statistically significant increase in FRH (p-value < 0.05) and much lower WC-1 levels (p-value = 0.0001) relative to the L-FRQ strain (Fig. 3d and Fig. S3a and d). A decrease in WC-1 was not observed in any other strain. Considering that the mutant WF pair strain has both a high amplitude of the *pfrq::luc* reporter (as is commonly seen in a FRQ-KO strain (Fig. 3b and Fig. S2b) and decreased WC-1 levels (Fig. 3d), this suggests that a mutation in the WF pair motif in the L-FRQ NTD is sufficient to either directly or indirectly destabilize WC-1. Taken together, these data indicate that the phospho-cluster and WF pair function as independent modules that influence circadian timing, with the WF pair differentially affecting core clock protein levels.

### The L-FRQ NTD WF pair impacts timekeeping in vivo in a temperature-dependent manner

With the demonstration that mutations in the L-FRQ NTD affected timekeeping, we next considered the context in which the L-FRQ NTD may have an important circadian regulatory function. As described above, splicing alters the ratio of S-FRQ to L-FRQ *in vivo* and, consequently, the amount of the L-FRQ NTD. This regulatory process enables the clock to properly respond to environmental temperature (9, 13). Thus, we hypothesized that the L-FRQ NTD mutants (i.e., the WF pair and the phospho-cluster) may be important determinants of the circadian temperature response. To examine this possibility, we replicated the luciferase assay performed on the NTD mutants above, but this time also investigated circadian period in these strains at 20 and 30 °C (Fig. 4a). We found that the L-FRQ^3pS/3A^ and L-FRQ strains had a similar trend in period increase, with the L-FRQ^3pS/3A^ strain increasing in period length by 2 to 3 hrs. at increasing temperatures, though the strains absolute periods were consistently different (Fig. 4a). This indicates that while temporal clock kinetics are altered in the phospho-cluster strain, the change is not temperature-dependent, making it unlikely that the FRQ NTD phospho-cluster plays a role in the clock’s response to temperature (Fig. 4a). However, we found that the L-FRQ^WF81AA^ strain was arrhythmic at 20 °C, with periods of 25.4 and 26.2 hours noted at 25 °C and 30 °C, respectively (Fig. 4a). This meaningfully different period trend/profile suggests that, indeed, the WF pair plays a role in maintaining robust rhythms in a temperature-dependent manner.

**Figure 4.**
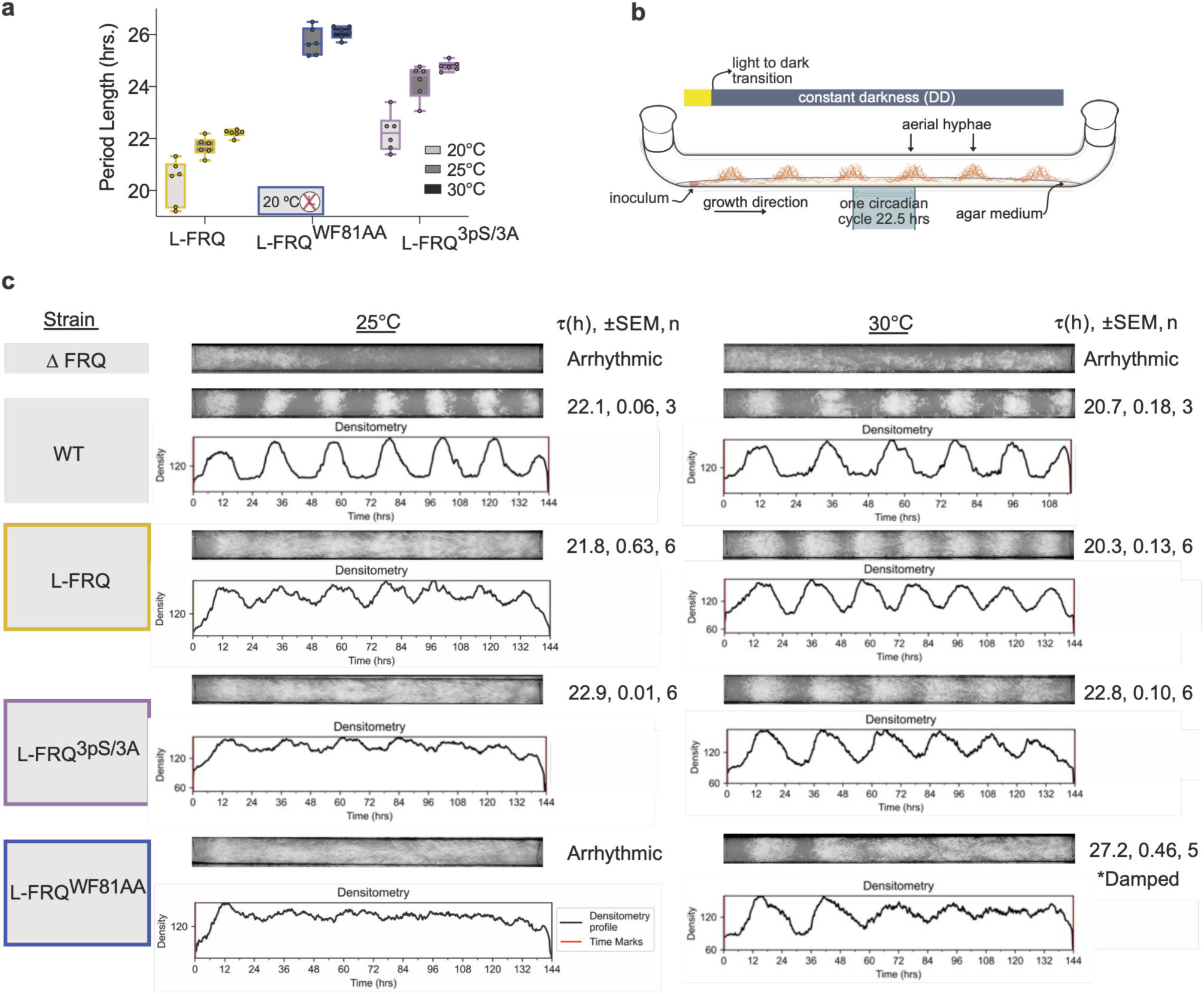
L-FRQ NTD mutants alter circadian periods and phenotypes across temperatures. **a**. Calculated periods for the L-FRQ NTD strains at 20, 25, and 30 °C as determined by luciferase analysis (n=6, fit with the Chronostar algorithm, see methods) (35). The horizontal bar in the box represents the mean, and the vertical bars represent the standard deviation. **b.** Cartoon schematic of a race tube assay adapted from (38). LL indicates constant light. **c.** Representative race tubes from race tube assays of the L-FRQ NTD strains at 25 and 30 °C.

Periods and densitometry plots were generated using Rhythmidia (38). Strain WT is a relative wild type and is *ras- 1^bd^* genotype. Periods were calculated using the Sokolov-Bushel periodogram (n=3 to 6; SEM is the standard error of the mean). Period calculation data are tabulated in Table S3.

To corroborate the effect of the WF pair in L-FRQ NTD on the maintenance of robust circadian rhythms over a range of physiological temperatures, we repeated the analysis of the above Neurospora strains in a race tube assay, which uses growth rate and tissue morphology changes under constant darkness and constant temperature as a proxy for phenotypic circadian period calculation (i.e., tracking the circadian output) (Fig. 4b) (38). Similar to the L-FRQ control, the L-FRQ^3pS/3A^ strain demonstrated persistent rhythms with a roughly equivalent period at 25 and 30°C, further suggesting that the phospho-cluster mutation does not impact the temperature response (Fig. 4c and Fig. S2c). Strikingly, there were no observable persistent phenotypic rhythms at 25°C in the L-FRQ^WF81AA^ strain (Fig. 4c and Fig. S2c). At warmer temperatures (30°C), the L-FRQ^WF81AA^ strain had a damped circadian oscillation with a 27.3-hour period. We were surprised by the notable discrepancy between our phenotypic race-tube data and our molecular circadian analysis. These differences are likely explained by the fact that banding measures output rather than the core clock oscillations, which is measured in the luciferase analysis, as output and core clock regulations are known to be governed differently(39). These data indicate that the WF pair in the L-FRQ NTD impacts timekeeping and clock stability in a temperature-dependent manner, directly affecting the output of circadian regulation, and that this is distinct from the role of the proximal phospho- cluster in circadian homeostasis (Fig. 3-4).

### The L-FRQ NTD encodes biochemical properties resulting in isoform-specific behavior

Since the L-FRQ NTD contains distinct motifs that influence temperature-dependent timekeeping, we asked whether the L-FRQ NTD itself confers regulatory potential distinct from that of the FRQ proteoform lacking the L-FRQ NTD, the Short FRQ isoform (S-FRQ). The difference between the isoforms is that S-FRQ lacks the L-FRQ NTD (a.a. 1-99, Fig. 5a). We therefore next constructed a Neurospora strain that expressed only the S-FRQ allele by altering the L-FRQ start codon to ensure that only S-FRQ is translated in the FRQ KO background (i.e., *pfrq::luc and ras-1^bd^*). Tracking luciferase levels in this strain over time across temperatures, we found that S-FRQ has a longer period at 25°C and 30°C than the L- FRQ and FRQ^D^ strains (Fig. 5b), consistent with previous observations (9, 13). Notably, the L-FRQ^3pS/3A^ but not the L-FRQ^WF81AA^ strain phenocopies the S-FRQ strain, suggesting that the phospho-cluster may be a key site that differentiates the phospho-kinetics of S-FRQ and L-FRQ (Figs. 4a and 5a).

**Figure 5.**
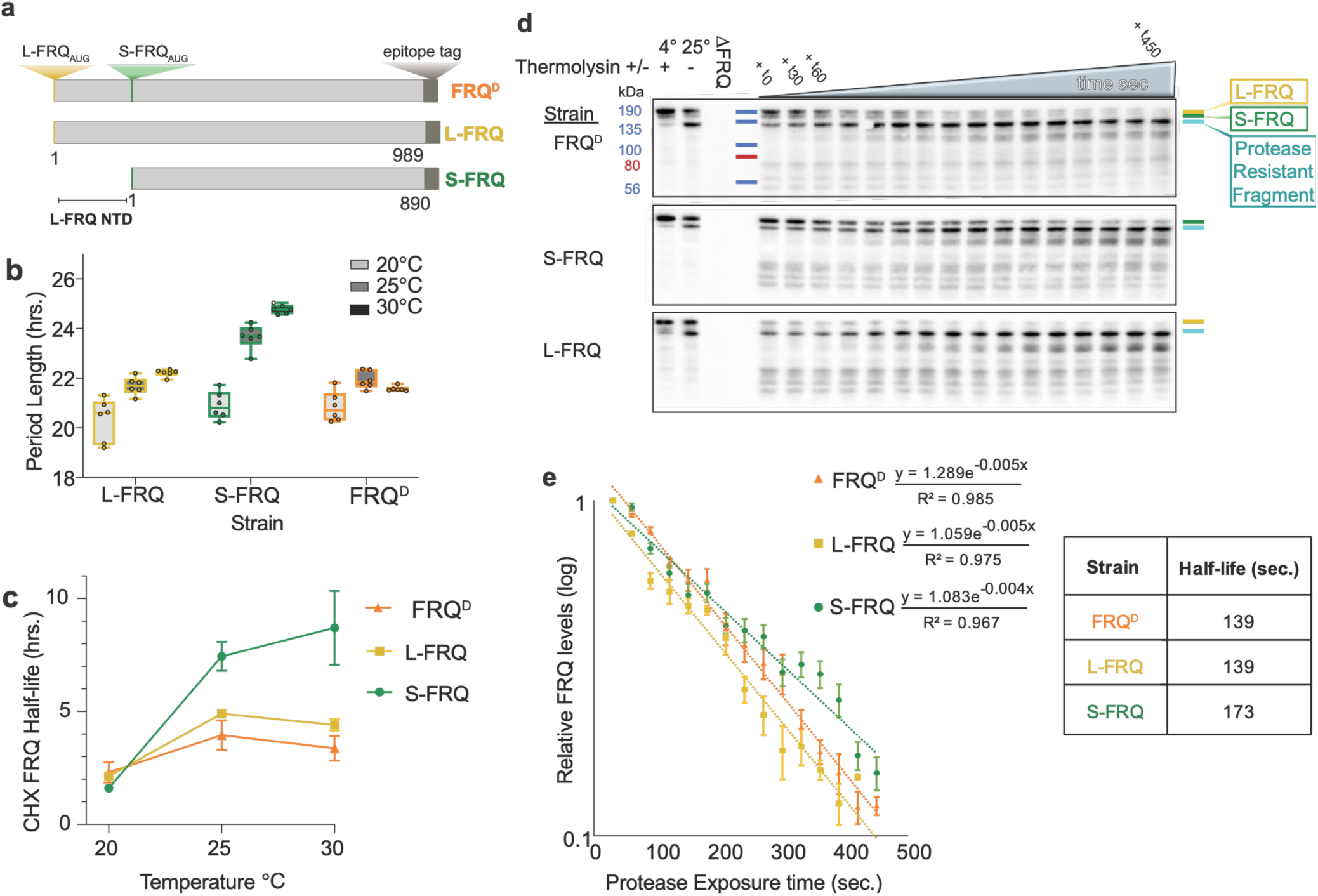
FRQ mono-isoform strains demonstrate isoform-specific correlations in period, half-life, and protease accessibility. **a**. Linear protein schematic representing the difference between the FRQ isoforms. Strain FRQ^D^ expresses both FRQ isoforms, while the S-FRQ and L-FRQ strains only express each respective isoform. **b.** Calculated periods for the FRQ mono-isoform strains and control as determined by luciferase analysis for replicates (n=6) at 20, 25, and 30 °C using the Chronostar algorithm over 96 hours (see methods). The control strain used is FRQ^D^, which expresses both isoforms. The horizontal bar in the box represents the mean, and the vertical bars represent the standard deviation. **c.** Protein half-lives calculated from a cycloheximide chase assay of cultures grown at 20, 25, and 30 °C. Points are means of n=3 replicates with SEM plotted at each point (note: the FRQ^D^ 25°C point is n=2). An exponential least-squares regression fit was used to calculate the t_1/2_ for each strain (data in Table S4) . **d.** Representative Western blots of the CRAFTY assay. Thermolysin was added at t_0_ at 25 °C, and the reaction was quenched at successive time intervals (t_30,_ t_60,_ etc. – t=seconds). The grey triangle represents increasing protease exposure time, with each lane being a new time point. Western blots were probed with anti-V5. **e.** Densitometry analysis of the relative levels of FRQ protein from the CRAFTY assay (n=3, SEM). In the table above, the FRQ protein was calculated using a decay fit, with the goodness of fit indicated by R².

To determine mechanistic differences in period and temperature regulation between L-FRQ and S-FRQ, we next investigated the overall stability of each FRQ isoform, as FRQ half-life stability has been suggested to tune period length. To do so, we treated mono-isoform and control strains with cycloheximide (CHX) to inhibit translation (40–42). We grew the S-FRQ, L-FRQ, and FRQ^D^ strains in constant light at 20°C, 25°C, and 30°C, added CHX, and measured changes in FRQ levels via western blot to determine FRQ half-life using first-order decay kinetics (Fig. 5c and Fig. S5) (43). All strains showed a non-monotonic increase in half-life with increasing temperature. The increase is notable, as reaction kinetics predict that half-life decreases with increasing temperature, suggesting that compensatory mechanisms are at play. As with the period, the half-life of the FRQ protein in the FRQ^D^ strain most closely mirrored that of the L-FRQ strain as temperature increased, likely because the dominant isotype is L-FRQ at higher temperatures (9). In accordance with its longer period, S-FRQ exhibited a longer half-life as the temperature increased. The discrepancy between the half-lives of S-FRQ and L-FRQ as temperatures rise, coupled with known temperature-dependent splicing (9, 40), indicates that the differential half-lives of the isoforms constitute a tunable mechanism by which the clock can facilitate period homeostasis across temperatures.

We next hypothesized that differences in solvent accessibility of the L-FRQ NTD could alter the interaction potential of L-FRQ and thus explain the differential behavior of L-FRQ (i.e., period and half-life). To test the solvent accessibility of the N-terminus of FRQ, we employed a limited proteolysis technique by co-opting our CiRcadian nAtive FasT parallel proteolYsis (CRAFTY) assay (21). CRAFTY uses protease accessibility as a proxy for IDR ensemble biases, since more compact IDRs are less protease-accessible, whereas more expanded regions are more protease-accessible. We leveraged the C-terminal epitope tag of our S-FRQ, L-FRQ, and FRQ^D^ strains to investigate differences in solvent accessibility between the L-FRQ NTD and the N-terminus of S-FRQ (44). We performed CRAFTY in triplicate on protein lysates extracted from the FRQ^D^, L-FRQ, and S-FRQ strains grown in constant light at 25°C and calculated changes in protease accessibility of FRQ protein via first-order decay kinetics (44) (see methods) (Fig. 5d-e and Fig. S6a-b).

When comparing the relative CRAFTY proteolysis rates of FRQ protein in the mono-isoform strains (S-FRQ and L-FRQ), we found that FRQ from the S-FRQ strain was more protease-resistant than FRQ from the L-FRQ strain (Fig. 5d and e). This suggests that relative to L-FRQ, the L-FRQ N-terminus is readily solvent-accessible. Additionally, our analysis demonstrated a rapid degradation of the FRQ protein into a C-terminal protease-resistant fragment of approximately 135 kDa in all three strains (Fig. 5d, blue line). Interestingly, the N-terminal fragment that is rapidly degraded in all strains spans the FRQ region, which, upon codon optimization, has been shown to yield a nonfunctional clock (45, 46). To determine whether any additional major degradation bands were present, we repeated the Western blot using the FRQ antibody. We found that the resistant band was consistently at the C-terminus (Fig. S6c).

### The L-FRQ NTD facilitates isoform-specific interactions, including those involving proteins that regulate temperature response

While ensemble changes and FRQ protein stability are obvious mechanisms for regulating circadian period, the molecular mechanisms by which environmental changes and L-FRQ NTD ensemble dynamics affect cellular control remain unclear. However, it is known that isoforms distinguished by disordered segments increase the complexity and capacity of cellular regulation by rewiring signaling networks through the modulation of isoform-specific protein-protein interactions (47–49). Moreover, our work has shown that FRQ has a highly complex, temporally dynamic interactome that may be modulated by conformational changes in FRQ (44). Therefore, we hypothesized that the presence of the isoform-specific region of L-FRQ (i.e., the L-FRQ NTD) could modulate the FRQ protein interactome, thereby imparting temperature-specific circadian regulation (26, 44, 50–55).

To validate this hypothesis, we investigated the interactomes of the S-FRQ and L-FRQ strains in triplicate under constant light conditions. Macromolecular complexes centered on FRQ were purified from each strain using a two-step Ni^+^ agarose/αFLAG co-immunoprecipitation process (44). Co-immunoprecipitated proteins were analyzed using Nano-Spray Liquid Chromatography-Mass Spectrometry/Mass Spectrometry (NS-LC-MS/MS) (see methods). We further refined this list by excluding proteins detected in samples from an untagged control strain (Ku-70a) or that did not meet the threshold for presence in two out of three triplicates in the respective mono-isoform strain. The detection threshold for the NS-LC-MS/MS results was determined empirically from an analysis of known FRQ-interacting proteins (see Methods), as we have previously done (44). A comparison of the interacting proteins revealed that S-FRQ and L-FRQ strains shared roughly 40 interacting partners (Fig. 6a, grey, center), including the established core clock interactors FRH and WC-1 (Table S5). However, beyond the shared interactors, we found 118 and 34 unique interactors in the L-FRQ and S-FRQ strains, respectively (Fig. 6A and Table S5), highlighting the increased interaction capacity of L-FRQ.

**Figure 6.**
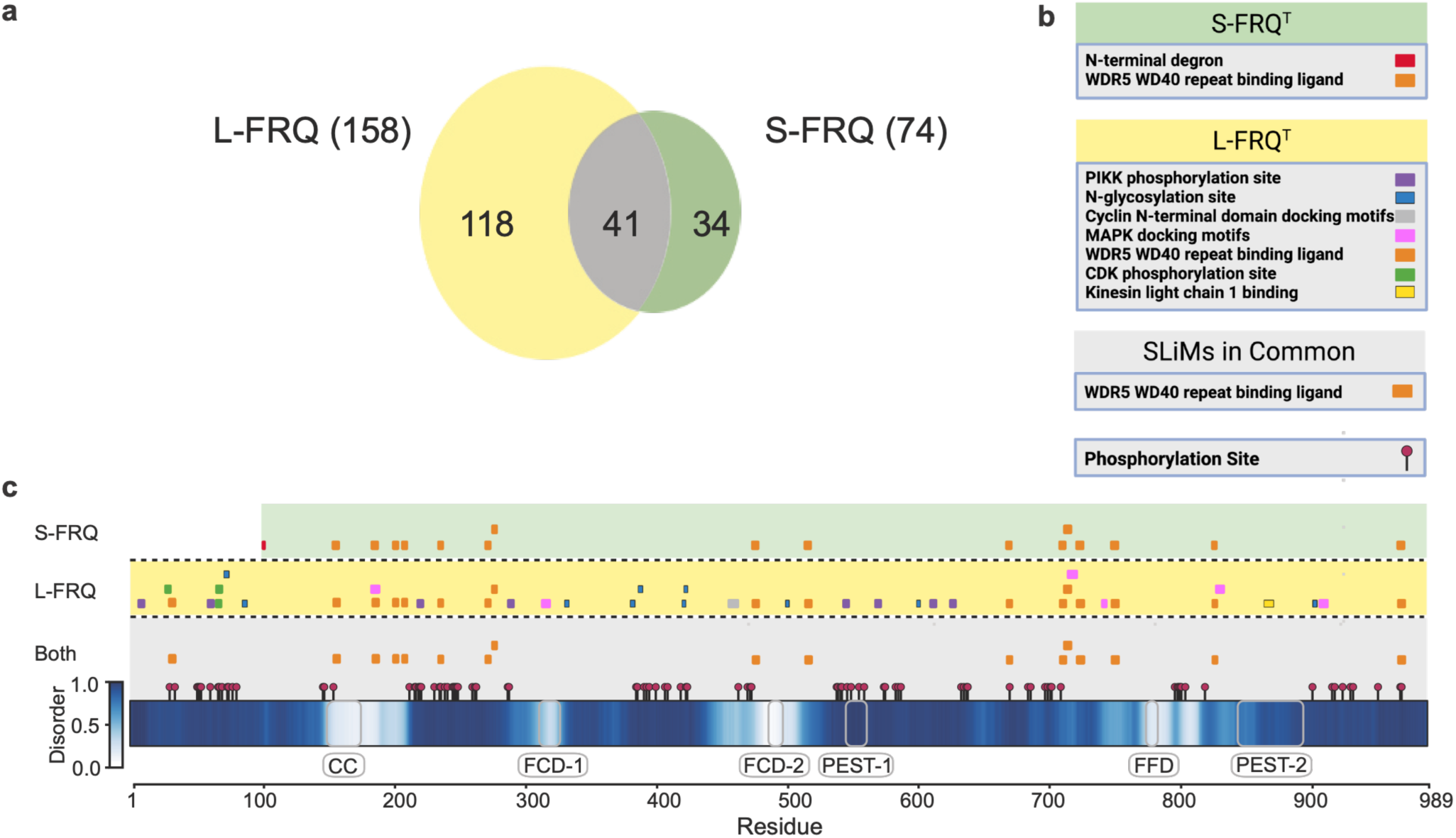
The FRQ isoforms have unique interacting partners, potentially facilitating differential circadian regulation. **a**. A Venn diagram illustrating the number of interacting proteins that are detected via Co-IP LC-MS/MS analysis in complex with FRQ in each of the mono-isoform strains (unique to L-FRQ (yellow), unique to S-FRQ only (green), and found in both datasets (gray)). **b.** Color-coded key of the different classifications of SLiMs predicted based on the interactions, grouped by the strain in which they were identified (60, 61). (**c.**) A FRQ protein diagram and alignment of predicted SLiMs from the mono-isiform strain interactomes. The known FRQ domains (grey-outlined boxes) and phospho-sites (maroon pins) are mapped to their locations (26, 44). The unique SLiMs with verified interactors for L-FRQ (yellow) and S-FRQ (green), and those found in common between the two datasets (gray). Below is a heatmap of predicted protein disorder levels. The hue corresponds with the predicted level of protein disorder across the sequence (24). Above 0.5, towards the darker hues, indicates the tendency towards disorder.

It has been shown that protein-protein interactions in IDRs can be mediated by Short Linear Motifs (SLiMs), 3-15-amino-acid sequences characterized by conserved residue patterns or residue properties (7, 56, 57). We and others have previously characterized the relationship among SLiMs, disorder, and FRQ phosphorylation, demonstrating a link between FRQ IDRs and protein binding (18, 44, 58, 59). Given these data, we hypothesized that, because each isoform has a unique interactome, we could identify FRQ isoform-specific SLiMs. To investigate the SLiM profile for each isoform, we performed an informatics analysis of the FRQ sequence for SLiMs using the Eukaryotic Linear Motif (ELM) finder (60, 61). We manually curated the SLiMs by excluding those for which the corresponding binder was not identified in the interactomes of S-FRQ, L-FRQ, or both. The verified SLiMs were mapped to a linear representation of the FRQ protein organized by each isoform (Fig. 6b and c). Furthermore, we annotated the known phosphorylation sites and disorder (26). Based on this approach, we observed correlations among disorder, interaction, and phosphorylation (Fig. 6b and c). We identified only one SLiM type unique to S-FRQ (i.e., the N-terminal degron) and six unique to L-FRQ. Given that L-FRQ has more unique SLiM types, it suggests that L-FRQ’s expanded interactome is at least in part facilitated by unique isoform-specific SLiMs in the disordered regions of FRQ (Fig. 6b and c).

We noted that L-FRQ was associated with a greater number of kinases (Table S5). In fact, the L-FRQ NTD housed the unique category “CDK phosphorylation site”, and SLiM distribution in the L-FRQ NTD aligns with the dynamically distinct regions identified in Figure 1 (Fig. 1f-g). Other L-FRQ-unique SLiMs across the remainder of the sequence include the “MAPK-docking motif” and “PIKK phosphorylation site” (Fig.6b). Importantly, MAPK (mitogen-activated protein kinase) pathways relay signals from environmental stressors throughout the cell and regulate the clock (62–66). Osmotic sensing-2, unique to the L-FRQ-only interactome (*os-2* or *hog-1*, NCU07024, Table S15), is a protein in the MAPK signaling cascade, which is known to bind the MAPK-docking SLiM. OS-2 undergoes FRQ-dependent circadian phosphorylation, which is important for both heat- and cold-stress responses (26, 62, 67–70). Taken together, these data suggest that the L-FRQ NTD rewires the FRQ interactome, facilitating interactions with known temperature-response pathways.

## Discussion

The molecular circadian clock is a paradigmatic biological system that depends on the molecular features of both structured domains and IDRs (7, 32, 71, 72). Although the structural regions of clock protein complexes are increasingly understood, much remains to be elucidated about the contributions of their disordered regions. In this study, we identified sequence-specific regulatory features of an IDR (i.e., L-FRQ NTD) that facilitate temperature-dependent tuning of clock repressors (Figs. 1-4). We examined a distinct region of the L-FRQ NTD (Fig. 1) and found that it exhibits ensemble features that enable context-specific regulation and modulation of molecular behavior (Figs. 2-4). Importantly, within the NTD, we identified two distinct regions that differentially influence the ensemble (Fig. 2). Mutating key residues within these regions altered clock function and robustness (Figs. 3-4). These findings are consistent with a growing body of work demonstrating that dynamic rewiring of IDRs via ensemble features and PTMs (i.e., phosphorylation) results in differential, tunable regulation (15, 73–76).

Another mechanism of molecular tuning associated with IDRs is alternative splicing. As previously mentioned, the L-FRQ NTD is alternatively spliced in a temperature-dependent manner (9, 13). Furthermore, temperature-regulated alternative splicing (AS) is widely used in animal and plant circadian timekeeping (7, 10, 12, 77, 78). Given that temperature-regulated AS in eukaryotic clock proteins yields multiple proteoforms with varying IDRs (7), this suggests that tunable isoform variation and the plastic nature of IDRs enhance timekeeping. Here, we have shown that in an alternatively spliced region, specific phospho-sites, in coordination with local chemistry, can affect the ensemble and, therefore, the molecular behavior of clock repressors (Figs. 2-5). Furthermore, our biophysical findings highlight that flexibility and disorder-associated features of alternatively spliced IDRs fine-tune period and expand clock regulation in a context-specific manner (Figs. 4-6).

Interactome rewiring resulting in increased biological regulation is a common feature of isoforms with alternatively spliced IDRs, and these features enhance post-transcriptional regulation (17, 48, 49, 79, 80). Our data indicate that alternatively spliced L-FRQ NTD enhances the regulatory capacity of the core clock in fungi (Figs. 5-6). Importantly, these findings can be extrapolated to eukaryotic clocks in animals, where multiple isoforms or paralogs of both repressor and activator proteins play key regulatory roles (7). Disorder-associated interactome rewiring could also underscore mechanisms underlying tissue specificity in animal clocks (44, 81–83). Taken together, our data highlight the emerging role of IDRs in facilitating both plasticity and robustness in biological systems. This work also underscores the importance of multiple proteoforms in enhancing and expanding clock timing and output.

We recently reported that disordered features of core clock repressors in animals and fungi can regulate the accessibility of protein-binding motifs and charge blocks, potentially conferring temporal post-transcriptional regulation (32, 44). Here, our work extends these findings by showing that FRQ isoforms have distinct interactomes and that these interactomes correlate with protein-binding motifs in disordered regions (Fig. 5-6). Furthermore, we found that L-FRQ has a larger interactome, suggesting that L-FRQ plays a distinct role in output regulation compared with S-FRQ (Fig. 6). Combined with the previous demonstration that L-FRQ has a more cytosolic distribution than S-FRQ (84), this suggests that, while both isoforms can regulate core clock timekeeping, L-FRQ may play an expanded role in cellular regulation.

At the molecular level, our findings regarding the dynamically distinct regions of the L-FRQ NTD have mechanistic implications for both the repressor and the activator proteins in the clock. In *N. crassa*, activator repression is mediated by FRQ and CKI, leading to hyperphosphorylation of WC-1 and WC-2 (Fig. 1a). We were surprised to observe WC-1 destabilization and a temperature-response phenotype upon mutation of the aromatic pair (WF81AA) in the L-FRQ NTD, but not upon mutation of the proximal phospho-cluster (S72A, S73A, S76A) (Fig. 3). Since aromatic groups in IDRs are known to facilitate interactions, our data suggest that the WF81AA mutations may cause loss of interaction with an important regulator, thereby driving the observed molecular and phenotypic changes. Given the phenotypes associated with the aromatic pair mutation (WF81AA), the L-FRQ aromatic pair may play a distinct role in regulating the repression of WC-1 via CKI. In this model, the mutated aromatic pair disrupts an interaction required for proper phospho-regulation of WC-1 by CKI and FRQ, thereby destabilizing WC- 1 and leading to loss of clock persistence (Figs. 3-4). Future work will uncover the mechanistic link between the WF81AA mutation and WC-1 destabilization.

### Limitations

Limitations of this work include the fact that although *N. crassa* is a syncytial organism, all our *ex vivo* biochemical methods are bulk assays. Therefore, we may be missing subtle nuances associated with isoform-specific proteoforms and the importance of spatial regulation using this *ex vivo* approach. It is also possible that multiple phospho-states of FRQ coordinate to achieve robust timekeeping, which we cannot capture here. Furthermore, our interactome data are from co-complexes isolated under constant light and thus do not capture FRQ interactions that may occur across the circadian cycle. Lastly, although we have predicted interaction motifs on FRQ, many of our identified interactors are likely indirect and result from scaffolding and complex formation.

### Conclusions

Here, we demonstrate that adding an IDR to a clock repressor protein in fungi via AS can modify and enhance clock function through temperature-regulated, dynamically distinct regions. Our findings support the model that the molecular clock capitalizes on multiple proteoforms (e.g., isoforms) to increase the clock’s robustness and regulatory reach. The results also suggest that these temperature-regulated, dynamically distinct regions may enhance functionality, enabling the tuning, flexibility, and context-dependent function of clock repressor proteins. Parallels with animal clocks are evident, given the extensive disorder, the presence of multiple proteoforms, and temperature-induced AS that yields different IDRs in repressor proteins from both kingdoms. These data will therefore enable the development of a model of how clock proteins coordinate to form the TTFL, which is essential for exploiting the clock to improve human health and treatment outcomes and to enhance precision in circadian medicine.

## Materials and Methods

### Recombinant L-FRQ NTD expression

The sequence coding for L-FRQ-NTD (residues M1 – I99) was inserted into a pET24a vector and expressed with a His_6_-SUMO tag at the N-terminus in M9 minimal medium containing ^15^N NH_4_Cl and ^13^C C_6_-glucose. Protein expression was induced with 0.5 mM IPTG at OD_600_ 0.5-0.8 at 20°C overnight under shaking at 160 rpm. Cells were harvested at 6000xg for 15 min at 4°C and pellet resuspended in Buffer A (20 mM Tris 150 mM NaCl 10 mM imidazole pH 7.4) added a pellet of protease inhibitor cocktail (Roche) and lysed with a cell disruptor (Constant Systems Ltd.) at 20 kpsi. The lysate was centrifuged at 20.000xg for 30 min at 4°C and the supernatant loaded onto a 5 ml HisTrap FF Ni-NTA resin column (GE Healthcare) preequilibrated in Buffer A and afterwards washed with 5 CV Buffer A. The protein was eluted using 15 ml Buffer A added 250 mM imidazole. Afterwards, 0.2 mg ULP protease was added to the eluate which was then dialyzed against 20 mM Tris 150 mM NaCl 1 mM DTT pH 7.4 overnight at 4°C. 0.1% (v/v) TFA was added to the dialysate and centrifuged at 20.000xg at 4°C for 10 min and then loaded onto a Zorbax C18 (Agilent) column preequilibrated in Buffer B (Milli-Q 0.1% (v/v) TFA) followed by 2 CV wash with Buffer B and eluted using a gradient 0% to 100% Buffer C (70% (v/v) acetonitrile 0.1% (v/v) TFA over 4 CV. The fractions containing pure double-labelled ^13^C, ^15^N –L-FRQ-NTD were afterwards lyophilized and finally resuspended in Buffer D (20 mM Na_2_HPO_4_ 150 mM NaCl pH 7.4).

### NMR and CD

All NMR samples contained either 1 mM or 0.750 mM ^13^C,^15^N-L-FRQ-NTD in 20 mM Na_2_HPO_4_, 150 mM NaCl, pH 7.4, 10% D_2_O, and 0.125 mM 4,4-dimethyl-4-silapentane-1-sulfonic acid (DSS). All spectra were acquired at 283.15 K on a Bruker AVANCE III 600, 750 MHz or 800 MHz (^1^H) spectrometers equipped with cryogenic probes. Free induction decays were processed using either NMRPipe or Topspin (Bruker Biospin) and subsequently analyzed in CcpNmr. DSS proton chemical shift was set to 0.00 ppm and used as an internal reference and heteronuclei were reference by relative gyromagnetic ratios. Assignment of backbone nuclei of FRQ-NTR were accomplished manually using ^15^N-^1^H-HSQC, HNCACB, HNCOCACB, HNCO, HNCACO, HNCANNH and HNCOCANNH spectra acquired using standard pulse sequences (Bruker BioSpin) and recorded with nonuniform sampling. and C^α^ SCSs of FRQ-NTR were calculated using random coil values. T_1_- and T_2_ ^15^N-relaxation times of ^15^N-FRQ-NTRs were acquired at 600 MHz (^1^H) on two series of ^15^N-HSQC spectra with varying relaxation delays (8 sepctra, 20-1.200 ms and 8 spectra, 0-271 ms) for T_1_ and T_2_, respectively including triplicate measurements. T_2_ relaxation data acquired at 10°C at 800 MHz (^1^H) using 8 (34-339 ms) different relaxation delays including triplicate measurements. All relaxation experiments were recorded using a 2 s recycle delay. Relaxation times were determined by fitting the relaxation decays to a single exponential in GraphPad Prism.

Far-UV CD spectra were recorded at 298 K on at 15 μM L-FRQ-NTD in 10 mM NaH2PO4, 50 mM NaF, pH 7.5 on a Jasco-J815. A total of 10 spectra recoded with a path length of 1 mm, ranging from 260-190 nm, data pitch of 0.1 nm and D.I.T of 2 sec, scanning spead of 20 nm/min. A corresponding CD spectrum of the buffer were recorded with identical setttings and substracted.

### Sequence analysis

Per-residue conservation across 12 related frq sequences taken from the *Sordariomycetes* class (*Neurospora crassa, Neurospora tetrasperma, Neurospora discrete, Sordaria macrospora, Fusarium graminearum, Fusarium verticillioides, Fusarium fujikuroi, Fusarium proliferatum, Fusarium oxysporum, Trichoderma reesei,* and *Trichoderma virens*) was calculated using the property-entropy metric of Mirny and Shakhnovich (85), as implemented in the score_conservation.py code of Capra and Singh(86). The alignment of these sequences was generated using Clustal Omega (ClustalW) with default settings (87).

Briefly, Henikoff sequence weights were computed across the multiple sequence alignment and used to build a weighted amino acid frequency distribution for each alignment column (with a 10^7^ pseudocount), and these frequencies were then summed into the six physicochemical classes as defined by Mirny and Shakhnovich ([AVLIMC], [FWYH], [STNQ], [KR], [DE], [GP]); the Shannon entropy of the resulting class distribution was normalized by log(min(n_classes, n_sequences)) and converted into a conservation score as 1 − H, which was multiplied by a gap penalty equal to one minus the weighted fraction of gaps in that column. The column-wise scores were projected back onto the ungapped positions of the *N. crasa* sequence, yielding one score per residue of that sequence, where higher values indicate stronger conservation of physicochemical character. Code for plotting this analysis is available at https://github.com/holehouse-lab/supportingdata/tree/master/2026/pelham_2026

### FINCHES DMS

Computational deep mutational scanning was performed using the dms() method implemented in the FINCHES frontend classes (introduced in FINCHES version 0.2.0). FINCHES enables prediction of IDR-mediated intermolecular interactions by repurposing chemical-physics principles in simple coarse-grained force fields as an analytical energy function (28). Here we introduce FINCHES-DMS, building on prior work to enable computational mutagenesis of key residues in an IDR.

For an input sequence of length *n*, every possible single amino acid substitution was constructed (20*n* variants), and for each variant the homotypic mean-field interaction parameter, epsilon, was calculated against itself using the Mpipi-GG force field(88, 89). Briefly, epsilon is obtained by building a pairwise interaction matrix over all residue pairs in the two sequences, in which each element is the finite integral of the pairwise potential (Wang–Frenkel plus Debye-screened Coulomb terms). This matrix is weighted by local sequence context to account for charge segregation and aliphatic clustering, partitioned into attractive and repulsive components at a poly-GS-calibrated null-interaction baseline, and reduced to a scalar by summing the per-residue mean attractive and repulsive contributions. This is the pipeline described in the original FINCHES paper, reproduced here for completeness.

Results from a FINCHES-DMS scan are assembled into a 20 × *n* matrix of Δε values, defined as the difference between variant and wild-type epsilon, such that negative values identify substitutions that increase self-interaction and positive values identify substitutions that decrease it. Because all variants are of equal length, values are directly comparable within a scan. The analysis considers single substitutions only and therefore does not capture epistasis. For more information, see the FINCHES documentation at https://finches.readthedocs.io.

### All-atom simulations

All-atom simulations of FRQ_1-99_ were performed using the ABSINTH implicit solvent model and the CAMPARI Monte Carlo (MC) simulation engine (https://campari.sourceforge.net/) (90). *N. crassa* FRQ_1-99_ plus an N-terminal Gly-Pro-Gly motif with N- and C-terminal caps (acetyl and amide groups, respectively) was simulated in a spherical droplet with radius = 149 Å. The monovalent ion concentration (NaCl) was set to 0.05 M and the temperature was set to 340 K to ensure adequate sampling of conformational space. The simulations presented each reflect a pool of ten independent replicates that commenced from a random coil starting conformation. Each simulation had 50 million steps following a 3.5 million step equilibration phase. The write-out frequency for accepted conformers was every 20,000 frames, which yielded a trajectory of 2375 frames per replicate (for 23,570 total frames). ABSINTH/OPLS-AA parameters were used for all simulations, including parameters for pSer from (30). WT FRQ_1-99,_ 3X- phosphorylated, WF>AA, and 3X-phosphorylated FRQ_1-99_ WF>AA sequences were all simulated using the settings and parameters described above.

### Simulations analyses

CAMPARI simulations were analyzed using MDTraj and SOURSOP using custom Python scripts (see Github) (91, 92). Ensemble distributions for the analytical Flory random coil (AFRC) model were generated using the AFRC Google Colab notebook (https://colab.research.google.com/drive/1WHw8ous7IgcKd2LKYuJLeBTlkdEYoRAk?usp=sharing) (93). Normalized distance maps were generated by calculating inter-residue distances from the simulated ensembles and dividing by the inter-residue distances from the AFRC model. Ensemble visualization was performed using VMD (94).

### FRET construct design, cloning and expression

The FRET backbone for bacterial expression (fIDP_pET-28a(+)-TEV) was prepared by ligating mTurquoise2 and mNeonGreen into pET28a-TEV or plasmid cloning DNA backbone using 5′ NdeI and 3′ XhoI restriction sites. Genes encoding for IDP regions were obtained from GenScript and ligated between the two FPs using 5′ SacI and 3′ HindIII restriction sites. Cloned plasmids were amplified in XL1 Blue cell lines (Thermo Fisher Scientific) using the manufacturer-supplied protocol. BL21(DE3) cells (Thermo Fisher Scientific) were transformed with fIDP_pET-28a(+)-TEV plasmids according to manufacturer protocol and grown in lysogeny broth medium with 50 μg ml^−1^ kanamycin. Cultures were incubated at 37 °C while shaking at 225 r.p.m. until optical density 600 of 0.6 was reached (approximately 3 h), then induced with 1 mM isopropyl β-d-1-thiogalactopyranoside and incubated for 20 h at 16 °C while shaking at 225 r.p.m. Cells were collected by centrifugation for 15 min at 3,000*g*, the supernatant was discarded and the cells were lysed in lysis buffer (50 mM NaH_2_PO_4_, pH 8 and 0.5 M NaCl) using a QSonica Q700 Sonicator (QSonica). Lysate was centrifuged for 1 h at 20,000*g* and the supernatant collected and flowed through a column packed with Ni-NTA beads (Qiagen). The FRET construct was eluted with 50 mM NaH_2_PO_4_, pH 8, 0.5 M NaCl and 250 mM imidazole, and further purified using size- exclusion chromatography on a Superdex 200 PG column (GE Healthcare) in an ÄKTA go protein purification system (GE Healthcare). The purified FRET constructs were divided into 200-μl aliquots, flash-frozen in liquid nitrogen and stored at −80 °C in 20 mM sodium phosphate buffer, pH 7.4, with the addition of 100 mM NaCl. Protein concentration was measured after thawing and before use using ultraviolet–visible (UV–vis) absorbance at 506 nm (the peak absorbance wavelength of mNeonGreen), and purity was assessed by sodium dodecyl-sulfate polyacrylamide gel electrophoresis after thawing and before use. To verify the brightness of the FPs, we measured the UV–vis absorbance of both donor (peak absorbance wavelength of 434 nm) and acceptor molecules before each FRET assay. We used only samples that displayed an absorbance ratio Abs_506_/Abs_434_ of 2.8 ± 0.2, a reasonable ratio given the difference in the molar extinction coefficients of mTurquoise2 and mNeonGreen (34,000 l mol cm^−2^ versus 116,000 l mol cm^−2^).

### In vitro FRET experiments and analysis

In vitro FRET experiments were conducted in black plastic 96-well plates (Nunc) with clear bottom using a CLARIOstar plate reader (BMG LABTECH). Buffer, stock solution and purified protein solution were mixed in each well to reach a volume of 150 μl containing the desired concentrations of the solute and the FRET construct, with a final concentration of 1 μM protein. Fluorescence measurements were taken from the top of the plate, at a focal height of 5.7 mm, with gain fixed at 1,020 for all samples. For each FRET construct, two repeats from different expressions with 6 or 12 technical replicates were performed in neat buffer (20 mM sodium phosphate buffer, pH 7.4, 100 mM NaCl), and two repeats from different expressions were done in every other solution condition. Fluorescence spectra were obtained for each FRET construct in each solution condition by exciting the sample in a 16-nm band centered at λ = 420 nm, with a dichroic at λ = 436.5 nm, and measuring fluorescence emission from λ = 450 to 600 nm, averaging over a 10-nm window moved at intervals of 0.5 nm. Base donor and acceptor spectra for each solution condition were obtained using the same excitation and emission parameters on solutions containing 1 μM mTurquoise2 or mNeonGreen alone. FRET efficiency calculations were performed as described previously(95).

### SEC and SAXS

Small-angle X-ray scattering (SAXS) experiments were performed at BioCAT (beamline 18ID at the Advanced Photon Source, Chicago). The experiments were performed with in-line size exclusion chromatography (SEC-SAXS) to separate monomeric protein from aggregates and improve the accuracy of buffer subtraction. Experiments were conducted at 22 °C in 20 mM sodium phosphate, pH 7.4, with 100 mM NaCl. Samples of approximately 300 µL were loaded, at a concentration of approximately 3.7 mg/mL, onto a Superdex 200 Increase 10/300 column (GE Life Sciences) and run at 0.6 mL/min using an ÄKTA Pure FPLC system (Cytiva). The column eluent passed through the UV monitor and proceeded through the SAXS flow cell, which consists of a 1.5 mm ID quartz capillary with 10 μm walls. The column to X-ray beam dead volume was approximately 0.1 mL. Scattering intensity was recorded using an Eiger2 XE 9M detector placed 3.685 m from the sample providing access to a q-range from 0.0029-0.42 Å-1. 0.5 second exposures were acquired every second during the elution. Data was reduced at the beamline using BioXTAS RAW version 2.1.4 (96, 97). The contribution of the buffer to the X-ray scattering curve was determined by averaging frames from the SEC eluent, which contained baseline levels of integrated X-ray scattering, UV absorbance, and conductance. Frames were selected as close to the protein elution as possible and, ideally, frames pre- and post-elution were averaged. Final scattering profiles were generated by subtracting the average buffer trace from all elution frames and averaging curves from elution volumes close to the maximum integrated scattering intensity; these frames were statistically similar in both small and large angles. Buffer subtraction and subsequent Guinier fits were done in BioXTAS RAW. Radii of gyration (Rg) were calculated from the slope of the fitted line of the Guinier plot at maximum qRg=1 using the equation(98):

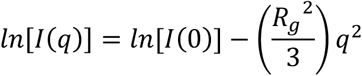

### Neurospora Strains

All FRQ mutant cassettes were recombineered for incorporation into the cyclophilin locus of Neurospora and transformed into strain uber#6 (*Δfrq::hph^+^, ras-1^bd^, his3::pfrq_c-box_-luc* ) or 122 (Δ*frq::hph*, *bd^+^*, mat a) strain as in (32, 44, 99). For the assembly template, genomic DNA was harvested from strain 74a, using the E.Z.N.A Fungal DNA Mini Kit from Omega (Cat: D3390-01). The L-FRQ cassette (*csr- 1::pfrq::lfrq^v5his103flag^*) had relevant start and stop codons modified by PCR, and a 10xGly, V5, 10xHis, and 3-FLAG epitope tag was fused to the C-terminus of the *frq* coding region, followed by a stop codon. The S-FRQ (*csr-1::pfrq::sfrq^v5his103flag+^*) and FRQ^D^ (*csr-1::pfrq::frq^v5his103flag^*) cassettes were assembled in the same manner. The L-FRQ NTD mutant strains (FRQ^WF81AA^ and FRQ^3pS/3A^) were generated by swapping the L-FRQ NTD from the L-FRQ cassette and incorporating a synthetic ultramer containing the relevant Ala substitutions. The cassettes were ligated using the NEBuilder kit (E5520S) and verified with automated entire construct sequencing.

Strains used as controls for race tube assays were 328-4 (*frq^+^, bd+, mat A*), uber #6 (*Δfrq::hph^+^, ras-1^bd^, his3::pfrq_c-box_-luc)* The negative control strain for the isoform mass spectrometric interactome analysis was Ku70a (*ku70::bar+)*.

### Period Analysis by Luciferase

Camera assays of *N. crassa* luminescence were conducted as in Jankowski et al 2023 (32). Low Nitrogen-CCD media (LN-CCD; 0.03% glucose, 0.05% arginine, 50 ng/mL biotin, 1xVogel’s salts, 1.5% bacto-agar, 25 µM luciferin, and 0.001 M Quinic Acid (pH 4.75)) was used for all luciferase reporter assays. 185 µL of this media was used per well in a white 96-well plate (Costar, 3912) and inoculated with 10 µL of the relevant conidial suspension. Plates were sealed with a breathable membrane (Breathe Easy, USA Scientific, 9123-6100) and incubated at 25°C in constant light (LL) for ∼48 hrs before being placed in continuous darkness (DD) in an incubator with a PIXIS CCD array (Princeton Instruments, 1024B) with a 35 mm Nikon DX lens (AF-S NIKKOR, 1:1 8G), run by the program Lightfield (version 5.2, Princeton Instruments). For the 20°C trial, plates were incubated at 25°C LL for ∼48 hrs to allow optimal growth, before being transferred to 20°C DD for image acquisition. The 25°C trial plate was grown at 25°C LL for ∼48 hrs. followed by 25°C DD, while the 30°C trial grew at 30°C LL for ∼48 hrs. followed by 30°C DD. Images were acquired for 15 min every hour, and final image stacks were imported into FIJI (ImageJ v2.0.0, NIH) to adjust brightness and remove noise using the default “Remove outlier” tool option. A custom image analysis plugin called “Toolset Image Analysis Larrondo’s Lab 1” (courtesy of Luis Larrondo) was used with its 96-well plate quantifying tool. From the background-corrected luminescence data, a custom Python script was used to determine the six wells most like each other for each strain and replicate. This was done over a portion of the experimental timeline by calculating the Euclidean distances between all the wells in the data series per strain for the span of 1-96 hours of the experiment and then using Ward’s method of hierarchical clustering (100) (script is available at https://github.com/Pelham-Lab). Once the six most similar wells were identified, a minimum threshold of two oscillations beyond the end of the entrainment period was used to classify each strain as rhythmic. Chronostar 3.0 was used to detrend the data using a 22-hour sliding window and the absolute space parameter (35). Periods were calculated from the fit of 6 replicates per strain and presented as a mean calculated value. The ΔFRQ KO strain was plotted as an average of 6 wells with the standard deviation.

### Culture Conditions and Protein Extraction for Constant Light Assays

Culturing *N. crassa* in constant light conditions was performed as in (21). Briefly, conidia were isolated via suspension and centrifugation from 5–7-day-old Vogel’s minimal slants of a given strain. The washed conidia were then inoculated into 50 mLs of Liquid Culture Media (LCM: 2% Glucose, 0.5% Arginine, 1xVogel’s Salts, 50ng/ml Biotin) and grown in constant light, with shaking at 125 rpm, for 48-72 hours. Samples were harvested using vacuum filtration, flash-frozen in liquid nitrogen, and stored at -80 °C until use. Protein was extracted from the mycelial mat by grinding the tissue in liquid nitrogen using a mortar and pestle and then resuspending the ground *N. crassa* in an equal volume of protein extraction buffer (PEB) (50 mM HEPES, 10% Glycerol, 0.4% NP-40 alternative,137 mM NaCl, 3mM CaCl_2_, pH 7.4, 1x HALT Protease and Phosphatase inhibitor ETDA free) to ground tissue. The ground tissue/PEB slurry was centrifuged at 14,000 RPM at 4 °C for 10 min, and the supernatant was removed and assessed for protein concentration with a Bradford assay.

### Lysate SDS-PAGE and Western blotting

Lysate protein concentrations were standardized, and samples were boiled with LDS Buffer (ThermoScientific, NP00007) and 2% v/v ß-mercaptoethanol for 5 min. Samples were then frozen at - 20°C until use. 13.8 µg of protein was loaded into each well of a precast NuPAGE 4-12% Bis-Tris gel (Invitrogen, WG1402BOX) in MES buffer for 50-80 min or NuPAGE 3-8% Tris-Acetate (Invitrogen WG1602BOX) in Tris-Acetate buffer for 60 minutes. The resultant gel was transferred according to the manufacturer’s specs using BioRad trans-blot turbo and BioRad RTA kits (Cat# 1704273). The membrane was blocked for one hour using 5% Milk in PBS buffer with 0.2% Tween-20. Dilutions of anti-V5 (1:5000, Cat# Invitrogen, 46-1157), immunodepleted anti-WC-1 (1:250, from (101)), and anti-FRH (1:10,000, from (102)) in PBS with 1% milk, 0.2% Tween were incubated overnight. Secondary concentrations were prepared by diluting 1:20,000 of Goat anti-Mouse (Invitrogen, 313430) or Goat anti-Rabbit (Invitrogen Cat# 31460) in PBS 0.2% Tween. SuperSignal West FEMTO (ThermoScientific, 34094) was used for chemiluminescent detection. All blots were normalized using amido black membrane staining. Imaging and relative quantification of Lysate Western blots was performed using Bio-Rad Image Lab software (version 6.0.1). Amido black staining of the membrane was used as a loading control for normalization. Data were plotted relative to the L-FRQ only strain in PRISM (version 10.1.1), and a one-way ANOVA was performed.

### Culture Conditions for Race Tube Period Analysis

Race tube assays were performed as in Keeley et al 2024, with slight modifications (38). Race tubes sourced from ChemGlass (cat. CG-4020-10) were filled with 15mL of race tube media (1×Vogel’s salts, 0.05% glucose, 0.1% arginine, 50 ng/mL biotin, 1.5% agar). Conidial suspensions were prepared from at least second-pass slants of desired strains with between 8.0x10^5^ and 1.5x10^6^ cells per tube. Race tube assays were conducted at 20°C, 25°C, and 30°C, respectively. Once inoculated, race tubes were incubated at their corresponding temperature first for 24 hours in constant light before being transferred to constant darkness for 6-7 further days and removed before reaching the end of the tube. Race tubes were marked once at the growth front upon transfer to darkness and once upon removal for analysis, yielding a true free run under constant conditions. Conditions of constant darkness and consistent temperature were confirmed for each experiment using Onset UA-002 pendant-style HOBO data loggers. Race tubes were scanned using an EPSON GT-1500 flatbed document scanner to produce a TIFF image of each pack of six tubes. These images were analyzed using Rhythmidia to determine circadian periods (38).

### Protein Stability Analysis

Cycloheximide protein half-life analysis was performed as described in (32) with minor adaptations. Conidial suspensions (see above) were inoculated into a petri dish containing 25mL LCM and grown in constant light at 20°C, 25°C, or 30°C for 24-26 hrs. Triplicate plugs were cut for each time point and cultured for 24-48 hours (strain dependent) in constant light in 50 mLs of LCM while shaking at 125 RPM. At timepoint 0, 40 µg/uL of Cycloheximide was added to all samples. Mycelial mats were harvested as above at hours 0, 2, 3, 4, 5, 6, and 7. Protein was then extracted as described above in lysate section. Lysate protein concentrations were standardized, and samples were boiled with 2x LDS Buffer (Cat #NP00007) 2% v/v ß-mercaptoethanol for 5 min. Samples were then frozen at -20°C until use. 13.8 µg of protein was loaded into each well of a precast NuPAGE 4-12% Bis-Tris gel (Invitrogen, WG1402BOX) in MES buffer for 50-80 min. The resultant gel was transferred according to the manufacturer’s specs using BioRad trans-blot turbo and BioRad RTA kits (1704273). The membrane was blocked for one hour using 5% Milk in PBS buffer with 0.2% Tween-20. A 1:5000 dilution of anti-V5 antibody in PBS with 1% milk, 0.2% Tween (Invitrogen, 46-1157), a 1:25000 dilution of Goat anti-Mouse (Invitrogen, 313430) in PBS 0.2% Tween, and SuperSignal West FEMTO (ThermoScientific, 34094) were used as the primary antibody, secondary antibody, and developer, respectively. All blots were normalized using amido black membrane staining. All steps of membrane processing, post-transfer and pre-substrate exposure, occurred in the automated Precision Biosystems Blotcycler (cat #W5WES1016BS) at 4°C. Quantification occurred as listed above in the previous western blotting section.

### CRAFTY Analysis

CRAFTY was adapted from (21, 44) to an extended protocol. Briefly, protein was extracted as described above. Lysate was standardized to 5 mg/mL and aliquoted on ice. Thermolysin solution (TL) (0.19 mM Thermolysin, pH 7.4, 50 mM HEPES, 137mM NaCl, 0.4% NP-40, 10% Glycerol) was added to the lysate at a ratio of 1 μL TL to 28.5 μL lysate. After mixing 20 μL/well of lysate with TL, the sample was aliquoted to a prechilled thermo cycler (with remaining sample left on ice). Once complete, the 96-well plate was heated to 25°C and the No TL sample was moved to a 25°C heat block. Immediately, 20 uL protein loading buffer (16.6, mM Tris–HCl pH 6.8, 0.66% SDS, 0.06% Bromophenol Blue, 3.3% Glycerol, 1% β- ME, 20 mM EDTA) was added to the first well. At each passing 30 second mark, 20uL was used to quench each successive well in the plate. This occurred over a period of 15 minutes. At the end of the digestion period, an equal volume (to that remaining) of protein loading buffer was added to the sample at 25°C with no TL, as well as the remaining aliquot on ice. All samples were then immediately heated to 95°C for 5 min, followed by a 4°C hold prior to storage in -20°C until SDS-PAGE. CRAFTY samples were western blotted in accordance with the protein stability analysis section above, with the exception of 15 ug of protein being loaded per well. Western-blotting with anti-V5, imaging, and quantification occurred as above.

### Protein Extraction and Purification for Mass Spectrometric Analyses of Neurospora

Interactomics was performed as in Pelham et al. (44). Tissue for NS-LC-MS/MS was generated by harvesting conidia from minimal media slants (50 mL size) by adding 50 mL of liquid culture media (LCM) and vortexing. Harvested conidia were resuspended in 4L of LCM, which was grown at 25°C in LL for 48 hours. *Neurospora* tissue (S-FRQ, L-FRQ, and Ku70a) was harvested by vacuum filtration and proteins were extracted using Protein Lysis Buffer with Halt^TM^ Protease and Phosphatase Inhibitor Cocktail diluted to a 1X concentration (Thermo Scientific^TM^ 78446). Protein concentration was measured through Bradford Assay and standardized to 35 mg/mL. 45 mL of total protein was transferred to washed/charged Ni-NTA agarose beads (Invitrogen^TM^ R90110) and incubated for 1 hour at 4°C with rotation before being washed. Proteins were eluted using The recommended Ni-NTA elution buffer. All buffers were prepared as described in the manufacturer’s instructions. The Ni-NTA eluted proteins were incubated with washed/pre-conjugated anti-FLAG magnetic beads (Millipore^TM^ M8823) overnight, washed, and eluted using Laemmle buffer. Beta-mercaptoethanol (BME) was added to an aliquot of protein elution for western blot analysis.

### Alkylation of Cysteine Residues

Proteins eluted from the anti-FLAG beads were reduced by adding 5mM of BME and incubated at 70°C for 20 minutes. Iodoacetamide was added to the sample to a final concentration of 13mM and incubated at room temperature in the dark for 1 hour. The alkylation was quenched by adding an additional 5mM of BME at 25°C for 15 minutes.

### Trichloroacetic Acid Precipitation and Protein Digestion

Alkylated protein products were mixed with cold 100% acetone and 100% trichloroacetic acid in a 1:8:1 ratio. This mixture was precipitated at -20°C for 1 hour before centrifugation at 11,500 rpm for 15 minutes at 4°C. The supernatant was discarded, and the protein pellet washed with 1 mL of cold 100% acetone and centrifuged again at 11,500 rpm for 15 minutes at 4°C. The supernatant was again discarded, and the protein pellet was air dried. The protein pellet was resuspended in 190 ul of 50 mM ammonium bicarbonate and digested with 20 ul of 1 ug/ul trypsin gold (Promega V5280) at 37°C for 4 hours.

### Analysis of FRQ Interactors by Mass Spectrometry

*N. crassa* protein pulldown samples were sent to the Mass-spec center at the University of Texas at Austin for analysis. Protein identification was performed via NS-LC-MS/MS using a Dionex Ultimate 3000 RSLCnano UPLC coupled to a Thermo Orbitrap Fusion. Prior to HPLC separation, the peptides were desalted using Millipore U-C18 ZipTip Pipette Tips following the manufacturer’’s protocol. A 2 cm long x 75 µm I.D. C18 trap column was followed by a 75 µm I.D. x 25 cm long analytical column packed with C18 3 µm material (Thermo Acclaim PepMap 100). Run-time was 1 hour. The FT-MS resolution was set to 120,000, and 3 sec cycle time MS/MS were acquired in ion trap mode. Raw data was processed using SEQUEST HT embedded in Proteome Discoverer. Scaffold (Proteome Software) was used for validation of peptide and protein identifications with filtering to achieve 99% protein confidence or a 1% FDR.

### Neurospora Interactome Data Analysis

The interactome was analyzed as described in Pelham et al., with slight modifications (44). To classify proteins as interactors (both in experimental and control conditions), protein thresholds in Scaffold were set at 99.9% and peptide thresholds were set at 95%, which are the minimum probability that a protein or peptide are identified in the spectra. Only proteins with ≥5% coverage or ≥4 unique peptides were considered as “identified” in our dataset. From the resultant Scaffold4 files, protein names, accession numbers, NCU numbers, number of peptides, and percent coverage for each protein (S-FRQ, L-FRQ, and Ku70a) were retrieved. The proteins identified in each S-FRQ and L-FRQ sample were retained if they were present in ≥5% total protein coverage or ≥4 unique peptides. The list was further refined by keeping only the proteins found in at least 2 out of the 3 replicates. In parallel, a master list of negative control proteins was generated of the proteins found in at least two replicates of the Ku70 samples using the same thresholds that were applied to the L-FRQ and S-FRQ protein lists. This master list of negative control proteins was then removed from the L and S-FRQ lists. The remaining NCUs of S-FRQ and L-FRQ lists were then compared yielding the overlapping “both” data set, as well as the independent S-FRQ and L-FRQ only datasets.

### Bioinformatic sequence analysis

The accession numbers for each interactome (S-FRQ, L-FRQ, and interactors common to both) were run through PANTHER (v19.0). The functional classifications and associated genes were exported and plotted. SLiM analysis was performed as in (44). The sequences of L-FRQ (NCU02265-t26_1) and S-FRQ (NCU02265-t26_2) were run through the eukaryotic linear motif predictor with a taxonomic context filter specific for the fungal kingdom using a motif probability cutoff of 100 (60). Since the ELM database has a high rate of false positives, the SLiMs were verified and retained through their protein interactors by manually cross-referencing with each interactome. The SLiMs were then mapped to a linear representation of FRQ in addition to the known phosphorylation sites from Baker et al 2009 (26). The SLiMs that are noted as occurring in “both” had proteins that occurred in both the S-FRQ strain and the L-FRQ strain pull-downs.

## Source data

Source data and code will be made publicly available upon publication if it is not already (see methods).

## Acknowledgements

This work was supported by the Washington University in St. Louis Department of Biochemistry and Molecular Biophysics Cori Fellow Program and Start-up (J.F.P.). This work was supported by the Novo Nordisk Foundation (#NNF18OC0033926; to B.B.K.). The NMR infrastructure at the Department of Biology, UCPH, was supported by the Novo Nordisk Foundation (#NNF18OC0032996 to B.B.K.) and Villum Fonden. A.S.H was supported by an NSF CAREER award (MCB-2338129). S.S. was funded by NIH- R35 GM137926 and acknowledges support from the Alfred P. Sloan Foundation. This work was supported by Rensselaer Polytechnic Institute Start-up funds and a grant from the NIH-NIGMS (R35GM128687) (J.M.H.). NIH-National Institute of General Medical Sciences T32 Fellowship GM067545 (to M.S.J.). E.T.U. was supported by NIH F32GM153018. We thank the members of the Pelham, Hurley, and Holehouse labs for support and suggestions.

## Classification

Major: Biological Sciences Minor: Biochemistry

