## Supplemental Information for "Proteoform plasticity modulates the temperature response in circadian timekeeping"

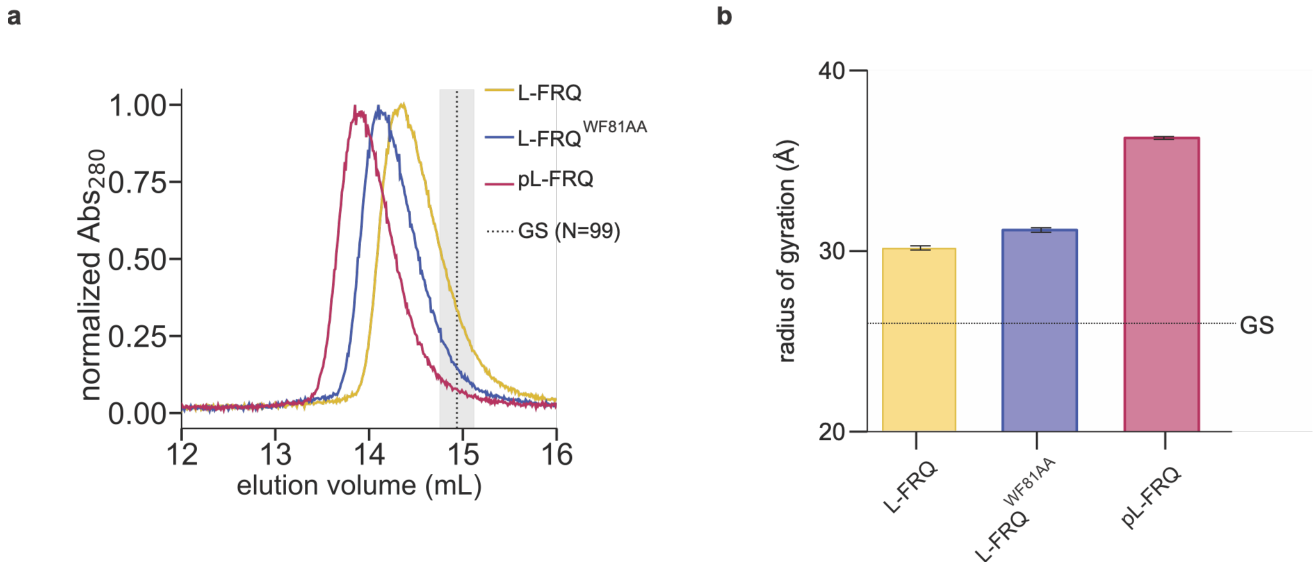


**Figure S1: L-FRQ NTD modification results in extended ensemble biases.** a. Size Exclusion Chromatography (SEC) profile of eFRET reporter constructs used in Fig 2. The dashed line represents the extrapolated elution volume of a glycine-serine (GS) repeat peptide control (1). b. Predicted radii of gyration of the L-FRQ NTD sequences calculated by all-atom Monte Carlo simulations. The dashed line is the Rg calculated for a GS repeat sequence of equivalent length.

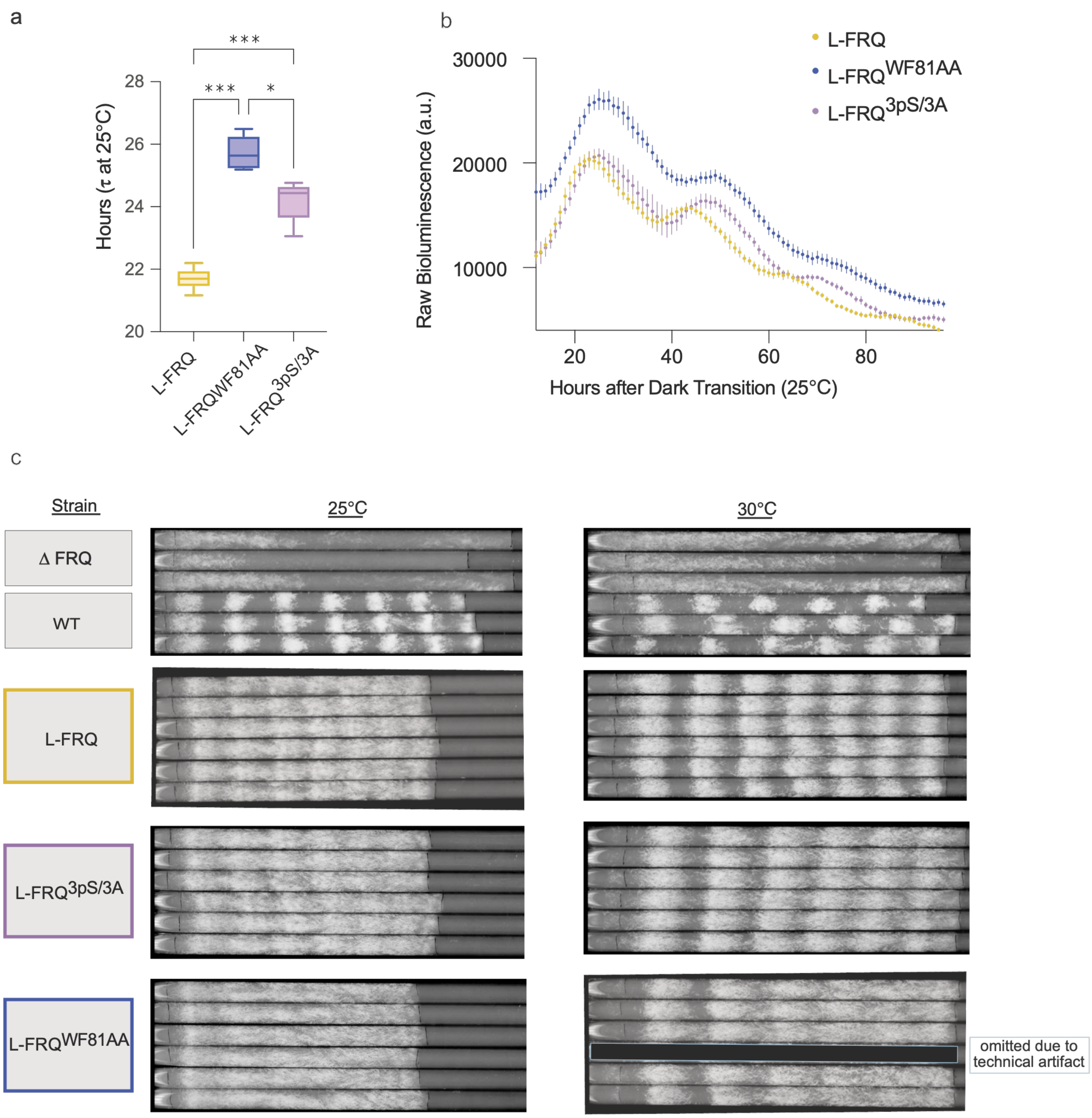


**Figure S2: L-FRQ NTD mutations result in an altered circadian period and race tube phenotypes.** a. Calculated periods for the L-FRQ NTD strains at 25°C determined by luciferase analysis (n=6, fit with the Chronostar algorithm, see methods) (2). Periods were tested with a one-way ANOVA (F = 70.15, p<0.0001) (***p<0.001, * p<0.05). b. Raw luminescence trace of L-FRQ NTD mutants. Points represent the average of n=6 wells, with bars indicating the standard deviation at each time point. An ordinary one-way ANOVA shows that the amplitude of the L-FRQ$^{WF81AA}$ strain is significantly different from the L-FRQ-only control strain, while the phospho-cluster strain is not (p<0.0001 and 0.5, respectively). c. Race tube images correspond to the data shown in Fig. 4. Race tubes were inoculated and cultured under constant light at their respective temperatures for 24 hours, then marked at the growth front and transferred to constant darkness at the same temperatures. Calculation data are provided in Table S3.

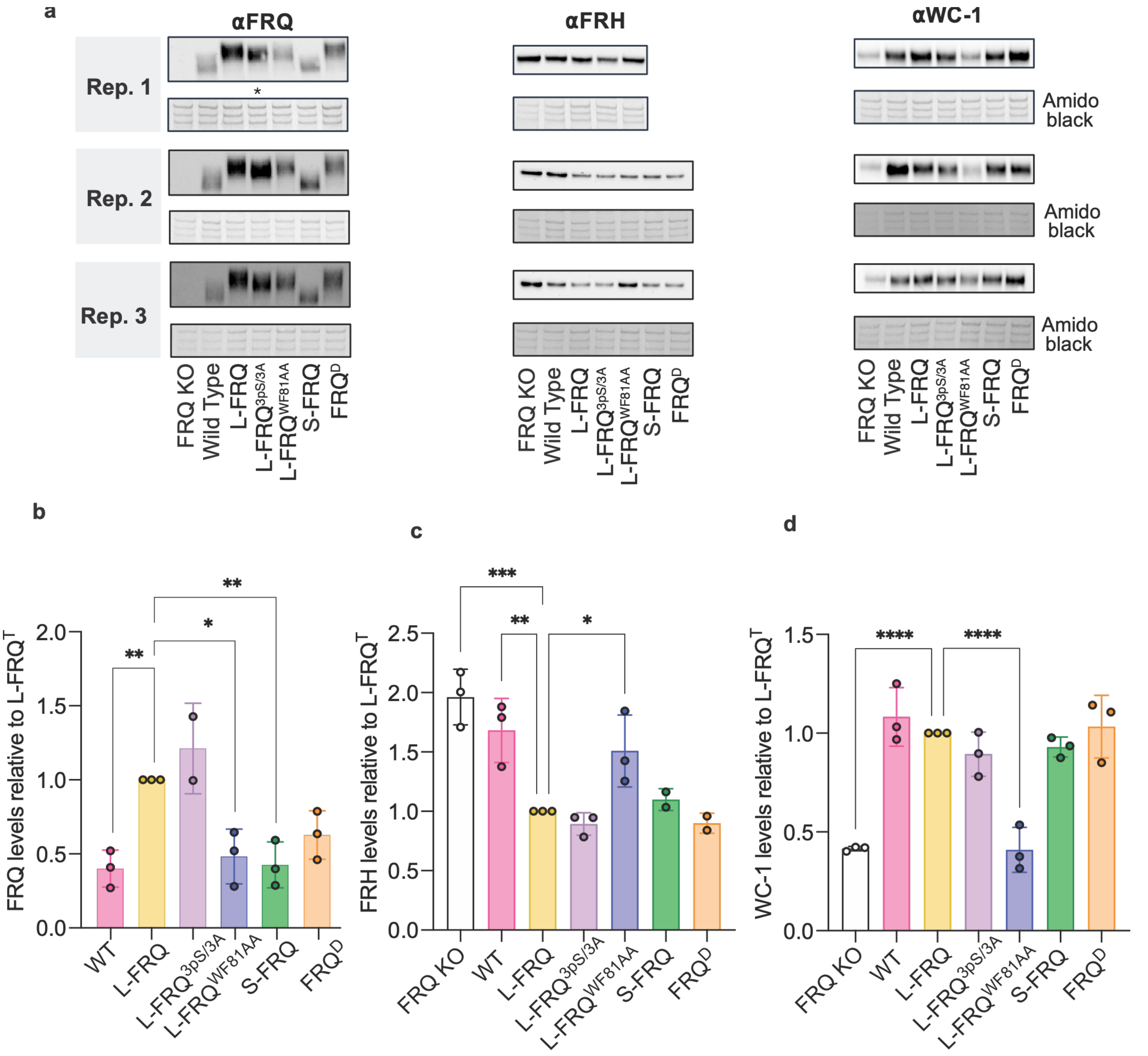


**Figure S3: Alterations of the FRQ NTD dynamically distinct regions result in changes in clock protein levels compared to the L-FRQ-only strain.** a. Western blots of clock proteins from FRQ mutant strains. The antibody used for probing is at the top of each column, and below each blot is the corresponding amido black stain used for normalization. A star indicates the lane was omitted from analysis due to an air bubble technical artifact. **b-d**. Relative quantification of the Western blots above is plotted relative to the L-FRQ strain for each clock protein. Amido black stains were used for total protein loading normalization. The statistical analysis used for each was a one-way ANOVA comparing values to L-FRQ, (**b**) FRQ, n=3 except where noted, F = 10.6, P = 0.0006, (**c**) FRH, n=3 except where noted, F = 12.90, P = 0.0001, and (**d**) n=3, WC-1 F = 23.09, P = <0.0001. ****p<0.0001, ***p<0.001, **p<0.01, *p<0.05. **d**. This is an extended version of the data presented in Fig. 3d.

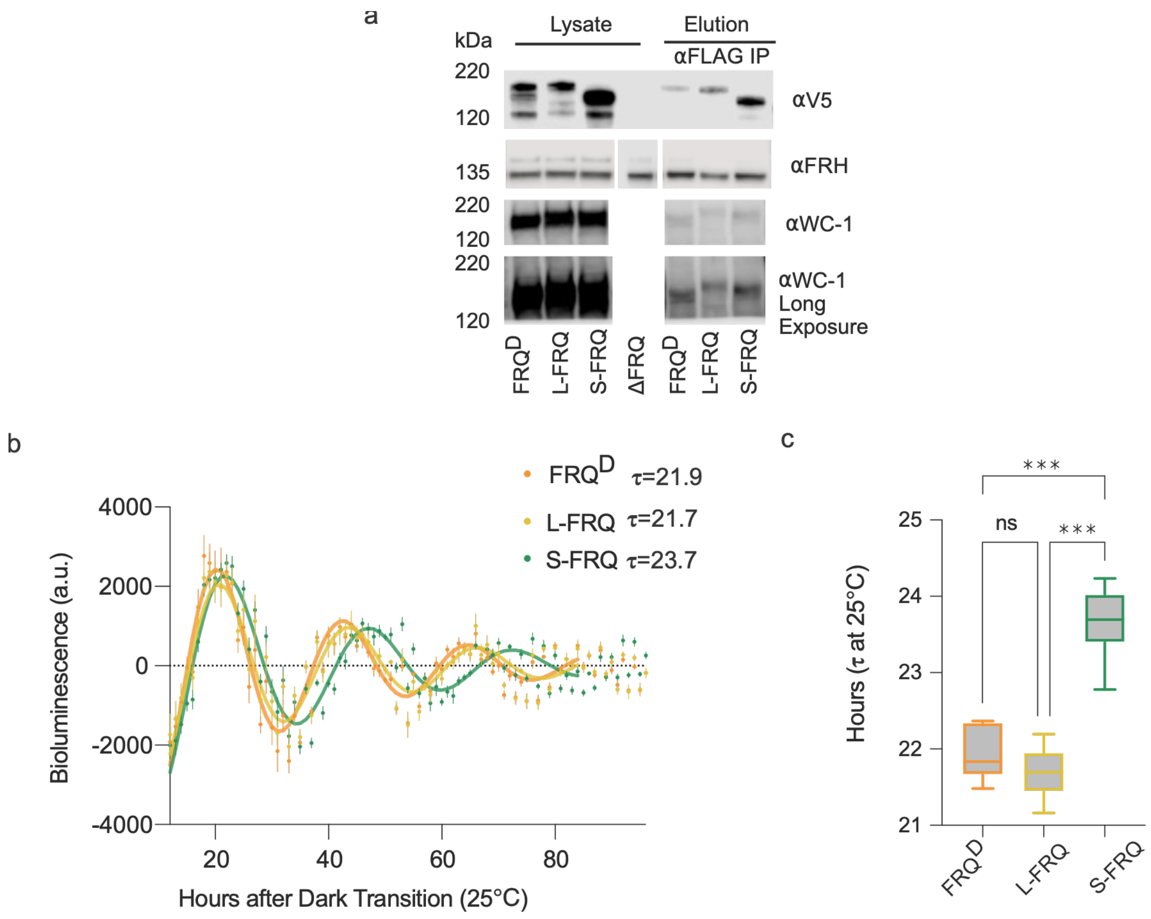


**Figure S4: Mono-isoform strains retain core clock interactions and rhythmicity**. **a.** Western blots of a Co-immunoprecipitation pulling down FRQ via the 3xFLAG epitope and probing for interactors FRH and WC-1. **b.** The smoothed, normalized, and detrended luciferase reporter activity of the L-FRQ, S-FRQ, and FRQ$^D$ strains. Points represent the average of n=6 wells, with bars indicating the standard deviation at each time point. The solid trace represents the fit determined by the Chronostar algorithm used to calculate the corresponding period ($\tau$) of each strain **c.** Calculated periods for the L-FRQ NTD strains at 25°C determined by luciferase analysis (n=6, fit with the Chronostar algorithm, see methods) (2). Periods were tested with a one-way ANOVA (F = 100.9, p<0.0001) (***p<0.001).

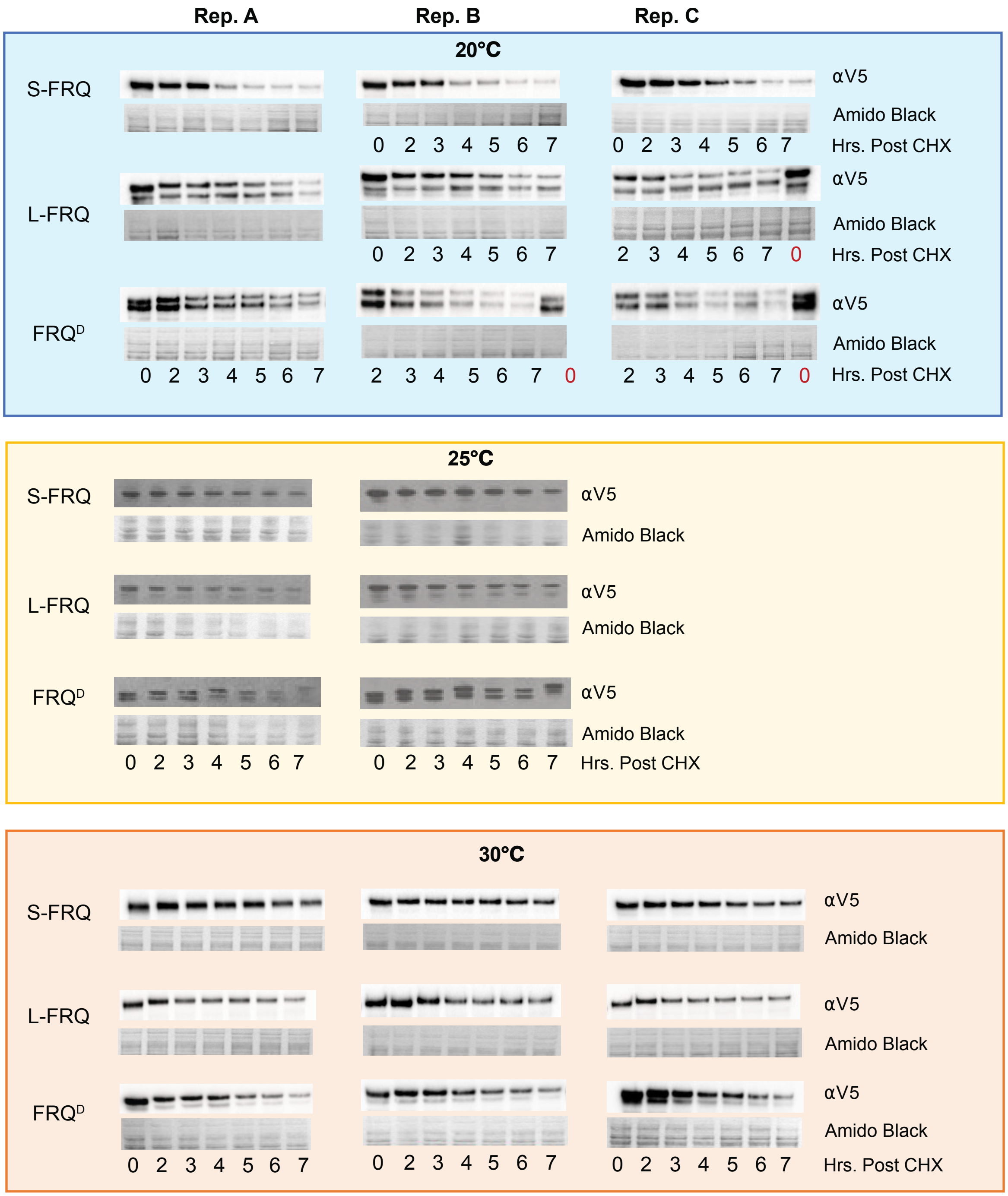


**Figure S5: Biological replicates of cycloheximide isoform analysis**. Western blots of the individual time courses of the CHX assay grouped by the three temperature conditions: 20°C (blue, top), 25°C (yellow, middle) and 30°C (orange, bottom) respectively. The red letter 0 indicates a different orientation of the samples compared to the other replicates. Calculations can be found in Table S4.

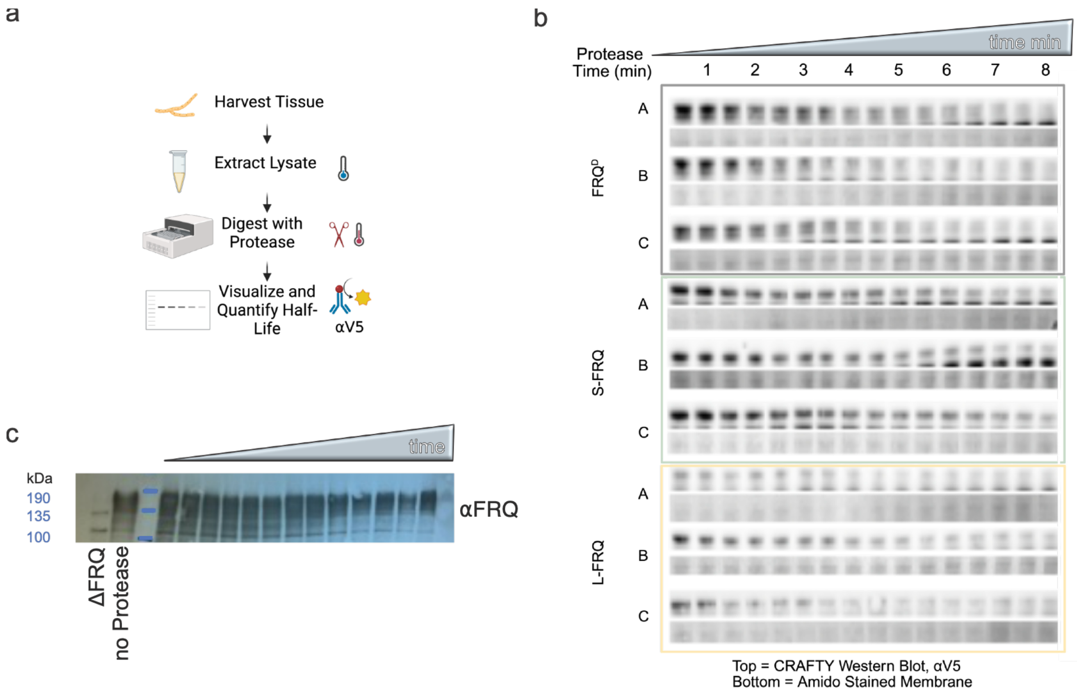


**Figure S6: Isoform strains have differential protease activity.** **a.** Schematic depicting the CRAFTY method (3). **b.** CRAFTY western blots used for quantification in Fig. 5d. Blots were probed with anti-V5. Below each blot is the corresponding amido black-stained membrane used for loading control normalization. **c.** CRAFTY western blot probed with anti-FRQ.
